# BZINB model-based pathway analysis and module identification facilitates integration of microbiome and metabolome data

**DOI:** 10.1101/2023.01.30.526301

**Authors:** Bridget Lin, Hunyong Cho, Chuwen Liu, Jeff Roach, Apoena Aguiar Ribeiro, Kimon Divaris, Di Wu

## Abstract

Integration of multi-omics data is a challenging but necessary step to advance our understanding of the biology underlying human health and disease processes. To date, investigations seeking to integrate multi-omics (e.g., microbiome and metabolome) employ simple correlation-based network analyses; however, these methods are not always well-suited for microbiome analyses because they do not accommodate the excess zeros typically present in these data. In this paper, we introduce a bivariate zero-inflated negative binomial (BZINB) model-based network and module analysis method that addresses this limitation and improves microbiome-metabolome correlation-based model fitting by accommodating excess zeros. We use real and simulated data based on a multi-omics study of childhood oral health (ZOE 2.0; investigating early childhood dental disease, ECC) and find that the accuracy of the BZINB model-based correlation method is superior compared to Spearman’s rank and Pearson correlations in terms of approximating the underlying relationships between microbial taxa and metabolites. The new method, BZINB-iMMPath facilitates the construction of metabolite-species and species-species correlation networks using BZINB and identifies modules of (i.e., correlated) species by combining BZINB and similarity-based clustering. Perturbations in correlation networks and modules can be efficiently tested between groups (i.e., healthy and diseased study participants). Upon application of the new method in the ZOE 2.0 study microbiome-metabolome data, we identify that several biologically-relevant correlations of ECC-associated microbial taxa with carbohydrate metabolites differ between healthy and dental caries-affected participants. In sum, we find that the BZINB model is a useful alternative to Spearman or Pearson correlations for estimating the underlying correlation of zero-inflated bivariate count data and thus is suitable for integrative analyses of multi-omics data such as those encountered in microbiome and metabolome studies.

## 1. Introduction

Microbiome data are essential for advancing our understanding of the biological basis of many human diseases and are becoming increasingly available. While descriptions of taxonomic aspects of the human microbiome are valuable, functional insights are arguably more informative. Accordingly, characterizations of the ways that bacteria interact with the host and the environment via metabolic byproducts and other biochemicals can offer important biological insights in disease pathogenesis, and offer targets for prevention and treatment. However, the complexity of these interactions cannot be underestimated. For example, relevant metabolites can be microbial products, whereas host-or environment-derived metabolites may serve as nutrients or environmental stressors for microbial communities. While the availability of microbiome-metabolome and health-disease associated phenotype data is increasing, suitable analysis methods development has not kept pace. Leveraging data on microbiome-metabolome interactions could help illuminate important biological pathways at play and identify bacterial species that influence each other via interspecies activities [1,2]. Importantly, these biological networks and microbial correlations may be influenced by the environment and differ between states of health and disease, as in the case of the oral biofilm microbiome-metabolome and dental caries [3,4]. Therefore, defining and measuring networks among microbial taxa, pathways in which taxa and metabolites are involved, and clusters of inter-correlated taxa are critical for understanding the function of microbial communities in health and diseases. Curated pathway datasets such as KEGG can provide known metabolic pathways involving metabolite networks but are not context-specific. The newly available Whole Genome Sequencing shotgun (WGS) DNAseq for metagenomics and RNAseq for metatranscriptomics (providing information at the at the taxon or gene level), or the earlier 16S sequencing for bacterial taxonomic abundance, paired with metabolome data from the same biofilm samples can provide unique new opportunities for context-specific integrative microbial pathway analyses.

Although joint network analyses of microbiome and metabolome data are critical for understanding host-microbiome interactions, the existing computational methods have not been designed for the specific characteristics of microbiome data. Until recently, Pearson or Spearman correlation-based pathway analyses [10] have been popular and robust for gene-gene network analysis for gene expression data; however, these approaches do not consider the excess zeros in microbiome data. Kendall’s Tau and Mutual Information (MI) have been suggested as possible replacements of Pearson or Spearman correlations for non-normal distributions, for example in single-cell RNAseq data [6–8]; however, MI is sensitive to threshold grids in data with excessive zeros, whereas Kendall’s Tau loses information on the continuous scale. More recently, copula-based pathway analysis [9] has been developed to model interactions between genes in single-cell RNAseq data while accommodating their non-normal distribution. Moreover, most existing approaches do not allow for between-group pathway change tests. Therefore, it is challenging to infer, for example, disease-specific microbiome-metabolome pathways and the essential hubs of microbial taxa and metabolites.

We propose a de novo pathway analysis that is independent of prior pathway knowledge and learns from the observed microbiome and metabolome data generated from matched samples (or at least from the same body sites or subjects, as long as a biological interaction hypothesis is valid). Our proposed method, BZINB-based integration of microbiome and metabolome for pathway analysis (BZINB-iMMPath), uses the newly developed bivariate zero inflated negative binomial (BZINB) model to directly model the joint distribution of a pair of count vectors, where one vector represents microbial species and the other vector represents metabolites, to estimate model-based correlations. The advantage of our method, which uses BZINB, is that we can rigorously handle the excess zeros in the distribution of microbiome counts [14]. Similar to single cell RNAseq data, microbiome data typically exhibit large numbers of zeros for several possible reasons, including the fact that some species may not be present in some samples, or structural zeros (e.g., due to technical artifacts, frequently referred to as “ dropout events”) represented by excess zeros in sequencing count data. Specifically, two advantages of using BZINB include the realistic assumptions of dropouts [15] in the zero inflated negative binomial (ZINB) distribution that allow the flexible modeling of both biological zeros (in the negative binomial component) and structural zeros (in logistic regression) to improve model fitting, and the feasibility of estimating correlations in the bivariate negative binomial (BNB) component conditional on the zero inflation component to reflect the underlying correlations.

We additionally propose, as another component of BZINB-iMMPath, the use of BZINB correlation measurements to represent the similarities [16] between species in species-wise clustering analysis to identify species modules (i.e., clusters) wherein species are highly correlated. Because the BZINB model accounts for zero inflation in a pair of species, or in individual species when investigating microbiome-metabolome correlations, most species and metabolites can be retained in the analysis rather than excluded because of zero-inflation, a feature that may be of biological importance.

To compare the accuracy of BZINB-based correlation with other popular correlation measures, we simulated pairs of correlated microbiome species and metabolite count vectors using the bivariate lognormal distribution and the BZINB distribution. We carried out simulations and applications using matched microbiome-metabolome data from a community-based study of childhood oral health/disease (ZOE 2.0 study, investigating early childhood caries or ECC) that sampled 3-5-year-old children’s supragingival dental biofilm. We also evaluated the accuracy of module identification using BZINB as a measure of similarity for cut-based clustering by crafting co-varying clusters of count vectors to represent species in semi-parametric simulations. We show that, in real data applications, the new method can identify the crafted clusters with high accuracy. Moreover, the integrated pathway analysis identified biologically significant and disease-specific microbial-metabolite pathways and meaningful inter-species interactions.

## 2. Materials and Methods

### 2.1. Description of BZINB Model

#### 2.1.1. ZINB model

Similar to single-cell data analysis, the probability of dropout per species per sample can be modeled using logistic regression in a framework of a zero-inflated model. The ZINB model has been previously proposed for the analysis of single-cell RNAseq data as a superior and more flexible model fitting compared to Poisson-based methods [13] for individual gene analyses in scRNAseq data, by allowing for both excess zeros and overdispersion.

#### 2.2.2. BNB model

Cho *et al*. 2021 began by introducing a bivariate negative binomial (BNB) model based on the Poisson-Gamma mixture model. First, let *R*_*j*_ *∼Gamma* (*α*_*j*_, *β*) for *j* = 0, 1, 2. Consider a pair of random variables (*X*_1_, *X*_2_), where *X*_1_ and *X*_2_ are each Poisson-distributed with means of *R*_0_ + *R*_1_ and *δ*(*R*_0_ + *R*_2_), respectively, where *δ* ∈ *R*^+^. These two mean variables are related through a common Gamma-distributed component, *R*_0_. Therefore, marginally, *X*_1_ and *X*_2_ each follow the negative binomial distribution such that 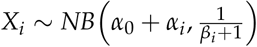 for *i* = 1, 2, where *β*_1_ = *β, β*_2_ = *δβ*. Thus, mean 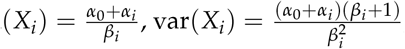, and 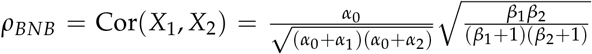. We henceforth denote (*X*_1_, *X*_2_) ∼ *BNB*(*α*_0_, *α*_1_, *α*_2_, *β*_1_, *β*_2_). Therefore, the parameters in *ρ*_*BNB*_ are estimated by fitting all the data to the BNB model.

#### 2.1.3. BZINB model

For correlation between a pair of genes in scRNaseq data, a bivariate zero-inflated (BZINB) model was proposed by Cho *et al*. 2021 that has the ZINB marginals, more parameters to flexibly accommodate the complexity of the single cell biology, and the estimated correlation conditional on the non-dropout events. With similar assumptions of dropouts observed as excess zeros and the overdispersion problem accentuated in microbiome data, here we extend the BZINB framework for microbial data modeling to compute a unique correlation measured between species or between species and metabolites. This new unique correlation analysis approach (i.e., BZINB-iMMPath) is model-based and uses the parameters estimated for the BNB component that is conditional on the probability of being non-dropouts in the BZINB model, defined as described below.

A pair of Bivariate Zero-Inflated Negative Binomial (BZINB) variables (*Y*_1_, *Y*_2_) ∼ *BZINB*(*α*_0_, *α*_1_, *α*_2_, *β*_1_, *β*_2_, *π*_1_, *π*_2_, *π*_3_, *π*_4_) follows a zero-inflated extension of the Bivariate Negative Binomial (BNB) distribution, where *π*_1_, *π*_2_, *π*_3_ and *π*_4_ respectively represent the probabilities of observing nonzero *Y*_1_ and *Y*_2_, nonzero *Y*_1_ only, nonzero *Y*_2_ only, and zero *Y*_1_ and *Y*_2_. Therefore, there is an underlying BNB component of the BZINB model, which is partially unobserved. Marginally, 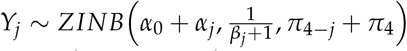 for *j* = 1, 2. In other words, without zeros, *Y*_*j*_ follows 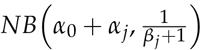, and each element of *Y*_*j*_ is zero with probability *π*_4_− _*j*_ + *π*_4_. Based on our understanding of excess zeros in the microbiome, the BNB components —which can include zeros from the negative binomial distribution— in the BZINB model reflect the underlying correlation between species after accounting for the dropouts (whether structural or technical) in BZINB. It follows that we use the same formula as *ρ*_*BNB*_ as in the model-based correlation. Therefore, we have 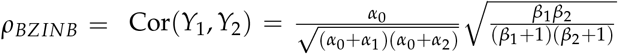 where all the parameters here are from the BNB component and are estimated by fitting all the data to the BZINB model. Although seemingly with the same format, the difference between our defined *ρ*_*BNB*_ and *ρ*_*BZINB*_ is whether we assume the presence of zero inflation in the data. Whether all of the data are used or not makes the two correlations different–this is due to the different assumptions in the models (BNB and BZINB) and the different meanings of (*α*_0_, …, *β*_2_) parameters between the two models.

There are variations of correlation in BZINB, such as the full-model BZINB correlation. That, besides the BNB component, also includes the zero component in the correlation. Simulation results (not shown) suggest this full BZINB model-based correlation introduces noise in the estimation and decreases the estimation accuracy of the underlying correlations.

### 2.2. Existing correlation calculation methods for network/pathway analysis

In correlation-based analysis such as network estimation for multi-omics count data, Pearson’s correlations are often used with the assumption of linearity. Previously, weighted correlation network analysis (WGCNA) has been used [10] to identify co-expressed clusters (modules) of highly correlated genes or other features. However, both microbiome and metabolome data contain excessive zeros, and therefore, there may be excessive ties in the data. In this case, Spearman’s rank correlation, even with less stringent assumptions compared to Pearson’s correlation, may still not be an appropriate measure.

In this study, we compare *ρ*_*BZINB*_ used in BZINB-iMMPath to not only *ρ*_*BNB*_ but also Spearman and Pearson correlation in terms of networks and module identification. The formula for Spearman correlation between vectors *X*_1_ = (*X*_1,1_, *X*_1,2_, …, *X*_1,*n*_) and *X*_2_ = (*X*_2,1_, *X*_2,2_, …, *X*_2,*n*_) is 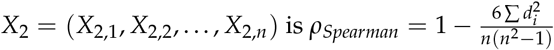, where *d*_*i*_ = rank(*X*_1*i*_) − rank(*X*_2,*i*_). In the case of ties, the average of the ranks is used. The formula for Pearson correlation is 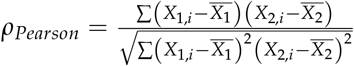.

### 2.3. Description of microbiome and metabolome data from the ZOE2.0 study

The ZOE2.0 study includes 6,404 3-5-year-old children enrolled in public preschools in North Carolina, United States, who underwent clinical dental examinations and biospecimen collection [23]. Of those, a subset of 300 participants’ supragingival biofilm samples were analyzed and made available for multi-omics (including metagenomics, metatranscriptomics, and metabolomics) analyses. Accordingly, 300 children have metagenomics data (WGS DNAseq, called DNA in this paper), 297 have metatranscriptomics (RNASeq) data, and 289 have metabolite data. Microbiome data have been made available via https://www.ncbi.nlm.nih.gov/bioproject/671299 and metabolome data via https://www.ebi.ac.uk/metabolights/MTBLS2215. As in a previous investigation (cite 33423574), ten participants with greater than 30% missing metabolite data and one ineligible participant were excluded. Among those 289 with metabolite data, 109 met the clinical criteria for ECC (i.e., cases) and 180 did not (i.e., non-cases). (cite: 30838597 and 30838598). To allow for comparisons of goodness-of-fit and variations in data sparsity (i.e., percentage of zeros) we used microbiome data generated by two different popular procedures for mapping and preprocessing metagenomics. Primarily, microbiome DNA data were classified into species-level profiles using a pipeline based on Kraken2 [17] and Bracken 2.5 [18] referred to as Kracken2/Bracken in this paper. The pipeline was built using a custom database including human, fungal, bacterial, and the expanded Human Oral Microbiome Database (eHOMD) [19] for microbial reference genomes. There were 417 microbial species identified as “ core species” after excluding rare and low-prevalence taxa that were kept in the analysis [5]. In a secondary procedure, the same DNA sequence reads were processed using MetaPhlAn2.2 through the HUMAnN 2.0 pipeline [11,12] with the default microbial reference genome in HUMAnN 2.0. Viruses, biosatellites, and unidentified species were filtered out, resulting in 205 species-level taxa remaining available for analysis. The advantage of Kraken2/Bracken for our application is due to the fact that it allowed the use of a custom and contemporary oral microbiome reference database and thus mapped oral/dental species more accurately than HUMAnN 2.0. On the other hand, HUMAnN 2.0 allowed not only the identification of species, but also the generation of gene-family and pathway-level data that can be of interest and value in some applications. The real data application of BZINB-iMMPath was done only using Kraken2/Bracken species-level data. Of note, all presented results rely on Kraken2/Bracken data unless HUMAnN 2.0 is explicitly mentioned, such as in goodness-of-fit and percentage of zeros comparisons that are presented in the Appendix.

The focus of the work reported in this paper is metagenomics data at the species level, but our new method can be applied to metatranscriptomics (i.e., RNAseq), as well as other levels of data, including gene-family or genes, because all data types are similarly characterized by excess zeros and overdispersion [20].

To obtain metabolomics data, samples were processed using Metabolon’s Ultra Performance Liquid Chromatography-tandem Mass Spectrometry pipeline [21,22]. A total of 503 named metabolites were identified through peak identification, QC, and correction for day-dependent technical variations [23]. Procedures and descriptions of the obtained metabolite data have been previously reported in detail (cite 33423574 and maybe even 34760716).

### 2.4. Simulation study

#### 2.4.1. lognormal based simulation

We simulated vectors representing pairs of metabolites and species, with theoretical correlations of 0.05, 0.1, 0.3, and 0.5, represent weak to strong correlations, based on the empirical distribution of correlations between the observed counts of pairs of species and metabolites (Figure 1). Each vector consisted of 300 elements drawn from a lognormal distribution, representing natural log-transformed counts. For simplicity, the marginal variance of the log-counts in each vector was set to 1, which was well in the range of the sample variances of the metabolite- and species-wise log-counts in ZOE2.0.

**Figure 1.**
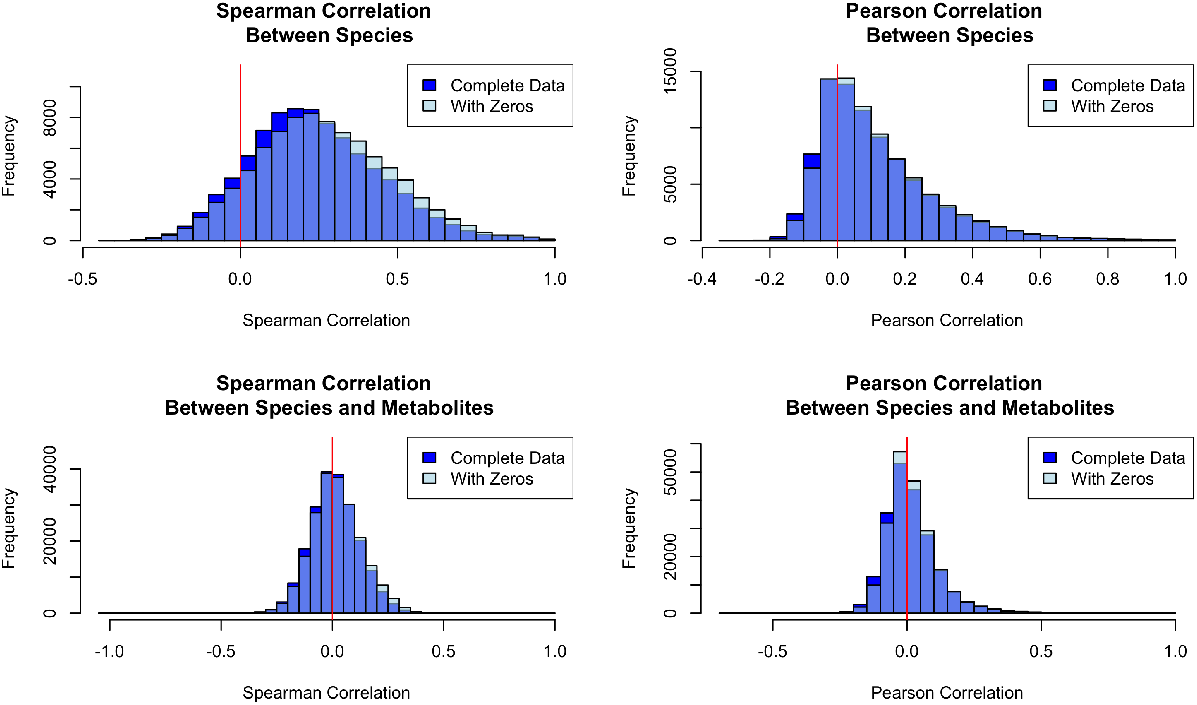
Empirical Spearman and Pearson correlations between pairs of (Kraken2/Bracken) species (417); and between pairs of (Kraken2/Bracken) species (417) and metabolites (503); in ZOE 2.0 (n=289). Correlations among complete data exclude subjects with one or more zeros in the pair, and correlations among data with zeros include all subjects.

Assuming that most missing values in metabolite data are due to low concentration, the counts in each metabolite vector were ranked and assigned a probability based on their rank. These probabilities spanned an interval of 0.3, centered at the pre-determined proportion missing. Let *rank*_*i*_ represent the rank of the *i*^*th*^ element in the metabolite vector, and let *p*_*zero*_ be the proportion of zeros in the vector. Then, the *i*^*th*^ element of the vector is set to zero with a probability of *p*_*i*_ = (0.5 − (*rank*_*i*_)/300) * 0.3 + *p*_*zero*_. Under the assumption that zeros in microbiome species are typically structural zeros, the elements in each vector representing a species were randomly chosen to be set to zero after the counts were simulated.

Figure 4 and Figure A3, illustrate the of number of zero counts against the mean of nonzero counts of each metabolite and species. These data revealed a decreasing trend in mean counts as the number of zeros increases and informed the selection of simulation parameters. Therefore, vector pairs representing metabolites and species were simulated under the scenarios outlined the first four rows in Table 1.

**Table 1.** Marginal log-scale means (before introducing zeros) and number of zeros for simulation of bivariate lognormal vectors with excess zeros that represent metabolite-species pairs (where *X*_1_ represents a metabolite, *X*_2_ represents a species) and species-species pairs. Levels of zero inflation include balanced (similar number of zeros in each vector) with either low or high zero inflation, and unbalanced (one vector has substantially more zeros than the other).

| Relationship |  | Zero Inflation | Number of zeros | Means |
| --- | --- | --- | --- | --- |
| Metabolite-Species | a | Balanced, low | 30, 60 | 14, 11 |
|  | b | Balanced, high | 150, 200 | 12, 9 |
| | c | $N_{zero,X_1} < N_{zero,X_2}$ | 30, 200 | 14, 9 |
| | d | $N_{zero,X_1} > N_{zero,X_2}$ | 150, 60 | 12, 11 |
| Species-Species | a | Balanced, low | 60, 60 | 11, 11 |
|  | b | Balanced, high | 200, 200 | 9, 9 |
| | c | $N_{zero,X_1} < N_{zero,X_2}$ | 60, 200 | 11, 9 |

In addition, the four correlation types were compared in simulated vector pairs that represent the relationships between two microbial species. These vectors were simulated based on the scenarios in the last three rows in Table 2. Zero counts were assigned randomly.

**Table 2.** Zero inflation parameters and resulting expected zeros for simulation of bivariate zeroinflated negative binomial vectors to represent pairs of metabolites and species (where *X*_1_ represents a metabolite, *X*_2_ represents a species) and pairs of species. Similar to the scenarios outlined in Table 1, there are balanced and unbalanced levels of zero inflation. The other BZINB model parameters are outlined in Supplementary Tables 1 and 2.

| | Zero Inflation | Expected zeros | $\pi_1$ | $\pi_2$ | $\pi_3$ | $\pi_4$ |
| --- | --- | --- | --- | --- | --- | --- |
| a | Balanced, low | 30, 60 | 0.75 | 0.15 | 0.05 | 0.05 |
| b | Balanced, high | 210, 240 | 0.1 | 0.2 | 0.1 | 0.6 |
| c | $N_{zero,X1} < N_{zero,X2}$ | 60, 225 | 0.2 | 0.6 | 0.05 | 0.15 |
| d | $N_{zero,X1} > N_{zero,X2}$ | 225, 60 | 0.2 | 0.05 | 0.6 | 0.15 |

#### 2.4.2. BZINB based simulation

To represent typical pairs as in the real data with various amounts of pairwise and non-pairwise zeros, vector pairs, we carried out a simulation using several combinations of parameters, as summarized in Table A1. For computational efficiency, these vector pairs represent rescaled pairs of count vectors obtained from the real data 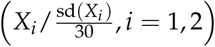, not altering the correlations. We considered underlying correlations of *ρ*_*BNB*_ = 0.05, 0.1, 0.30, and 0 by using different combinations of shape and scale parameters in the BZINB distribution (Table A1). For each combination of shape and scale parameters (and accordingly, level of correlation of the nonzero counts), we conducted simulations using four combinations of zero inflation parameters (*π*_1_, *π*_2_, *π*_3_, *π*_4_), representing balanced low, unbalanced, and balanced high zero inflations (Table 2).

We also simulated vector pairs under the BZINB distribution to represent typical pairs of microbial species. These vectors had the same zero inflation parameters as the microbiome-metabolome simulated vector pairs (Table 2), but different means and slightly different correlations. The corresponding shape and scale parameters are presented in Table A2.

### 2.5. Spectral clustering for module identification

#### 2.5.1. Approach for BZINB application in spectral clustering

Spectral clustering is a flexible method for partitioning networks using the eigenvectors of nodes’ similarity matrices [16], and has been used in many applications, including bioinformatics. Although similarity is typically quantified through the Gaussian kernel, other measures such as cosine similarity [24] have been used to better represent certain data types. In correlation networks, the positive correlation between a pair of nodes (or, in our data, species or metabolites) is scale-invariant and is often used as a measure of similarity when the co-varying dynamics of the nodes is of interest. Therefore, one can reasonably use the estimated correlations in constructing affinity matrices in applications such as spectral clustering to discover novel pathways that differ between study groups or potentially associated with health or disease states. In this paper, we compare the Spearman, BNB, and BZINB correlations in spectral clustering for microbiome count data.

For vectors *x*_*i*_ and *x*_*j*_, the affinity *a*_*ij*_ is a measure of similarity such that *a*_*ij*_ is bounded by 0 and 1, *a*_*ij*_ is closer to 1 as *x*_*i*_ and *x*_*j*_ are more similar, and *a*_*i*_ *j* = 0 when *i* = *j*. To obtain each affinity matrix from a correlation matrix, we set the diagonal entries to zero. Since the BZINB model-based correlation can only be positive, we force any negative values obtained from Spearman correlations to be zero. This allows us to only predict clusters with and based on positive inter-dependencies. Next, we cluster the nodes using SpectraLib_A [25]. While the affinity matrices are all symmetric, this method can account for directed networks, for example, to incorporate known interactions between species or metabolites, by using asymmetric affinity matrices.

#### 2.5.2. Evaluation of cut-based spectral clustering using crafted semi-parametric simulation

We simulated correlated clusters to compare the accuracy of the three types of affinity matrices as follows. We permuted the first 400 species in the caries-free (i.e., healthy-group) ZOE 2.0 participants and split them into 10 clusters of 40 species each. For each cluster k, we generated a random vector *R*_*k*_ ∼ *Pois*(17, 968) (since 17,968 was the mean count of the 400 species). For the nonzero counts of each species j in cluster k, we computed a weighted sum, *Z*_*j*_ = 0.9 * *Y*_*j*_ + 0.1 * *R*_*k*_, of each original species’ counts (*ϒ*_*j*_) and the random vector, to introduce additional correlation within each cluster. We then estimated the Spearman, BNB, and BZINB correlations between all 400 species to construct three types of affinity matrices. Then, we clustered the species, for each affinity matrix, using SpectraLib_A with 10 clusters. In cases where biological knowledge exists regarding the direction of effects in relationships between different ‘omics layers, the affinity matrix can be altered to reflect it.

To evaluate the accuracy of each correlation type in spectral clustering, we contrasted predicted and assigned clusters to optimize the prediction accuracy as follows:

1. If the most common predicted cluster for an assigned cluster is the same as the most common assigned cluster for that predicted cluster, those clusters are matched.
2. Then, the overall proportion of accurate predicted cluster assignments is calculated for each possible combination of the remaining clusters.
3. The remaining clusters are matched with the combination that maximizes the proportion of accurate predicted cluster assignments.

### 2.6. Network visualization

To create visual representations of networks, we represented each metabolite and each species as a node, and each correlation as an edge. For easier interpretation of the network diagrams, we included only a subset of metabolites and species. Heimisdottir *et al*. 2021 identified 16 metabolites and Cho *et al*. 2022 identified 15 species in ZOE 2.0 that were significantly associated with the childhood dental disease outcome of interest (i.e., ECC). In this work, we are focus on the patterns of co-occurrence between these species and metabolites, and examine whether these patterns differ between health and disease states. In network visualizations, we include only the strongest correlations, which are of interest. We visually assessed histograms of all correlations for each correlation type and disease group to determine optimal correlation cutoff points. We applied Cytoscape’s Organic layout and removed node overlaps. To accomplish this, we first obtained the BZINB-based and Spearman correlations between each pair of 16 metabolites and 15 species of interest, as well as between each pair of the 15 species in from ZOE2.0, in each of the two heath/disease (non-ECC/ECC) participant groups. Next, we sought to determine optimal cutoff correlation values to prevent the network visualization from being too large, even for 16 metabolites and 15 species. Therefore, we created network visualizations only for the most correlated species and the most correlated species-metabolites for the ECC and the non-ECC groups. To maintain comparability of the network diagrams, we used the same percentage of strongest correlations for each. After comparing several cut-off values, we determined that using the top 30% of metabolite-species correlations resulted in approximately 100 edges when the two disease groups were plotted on the same diagram, so that the edges and nodes were mostly visible while the network was large enough to illustrate high-degree nodes.

Network visualizations were generated with Cytoscape 3.9.1 [28]. Metabolite superpathways were highlighted by node color, and edge stroke color was used to denote health/disease (non-ECC/ECC) when correlations from both participant groups were plotted together.

## 3. Results

### 3.1. The BZINB model is a good fit for the ZOE2.0 microbiome and metabolome data

First, we sought to identify suitable distributions to model the paired metabolome and species-level microbiome count data. We assumed that proper normalization in microbiome and metabolome data have been carried out. Zeros present in the original counts will remain as zeros after normalization (RPK, RPKM or CPM).

Specifically, we evaluated model fits of three distributions with multiple randomly selected pairs of species and metabolites from ZOE2.0. Count data naturally correspond to a Poisson distribution, while the negative binomial distribution is an extension of Poisson that allows for overdispersion. Non-zero data can be transformed to lognormal to improve fit, particularly due to the long right-tailed distribution. It is important to consider that many species and metabolites exhibit large proportions of zeros. Therefore, candidate distributions included (1) zero-inflated Poisson, (2) zero-inflated negative binomial, and (3) zero-inflated lognormal. For each vector, model parameters were estimated using the nonzero counts from the real data. Numbers of zeros were simulated following a binomial distribution with probability p equal to the proportion of zeros in the real-data vector, and the remaining counts were simulated based on the estimated model parameters.

The simulated vectors from the zero-inflated Poisson distribution did not capture the overdispersion in most of the real-data vectors (Figure A1). The zero-inflated negative binomial distribution was found to adequately capture the data distribution of metabolite and microbiome (Figure 2). Because the negative binomial distribution takes on discrete values, we did not evaluate goodness-of-fit using the Kolmogorov-Smirnov test in this case. Further, using the Kolmogorov-Smirnov test, we assessed the goodness-of-fit of the lognormal distribution to metabolite and species data in ZOE2.0 (Figure 3). Because the Kolmogorov-Smirnov test is only applicable to continuous distributions, only the nonzero counts were included. 11.5% of metabolites had p-values less than 0.05, suggesting that the zero-inflated lognormal distribution was a good fit for most metabolite data. On the other hand, the zero-inflated lognormal distribution was not a good fit for over 20% of the Kraken2/Bracken species while it was a good fit for almost all HUMAnN 2.0-derived species in ZOE2.0 (Figure A2). Also, based on a visual comparison of Kraken2/Bracken real data and simulated zero-inflated lognormal count vectors (Figure 2), the zero-inflated lognormal distribution appeared to represent species data well.

**Figure 2.**
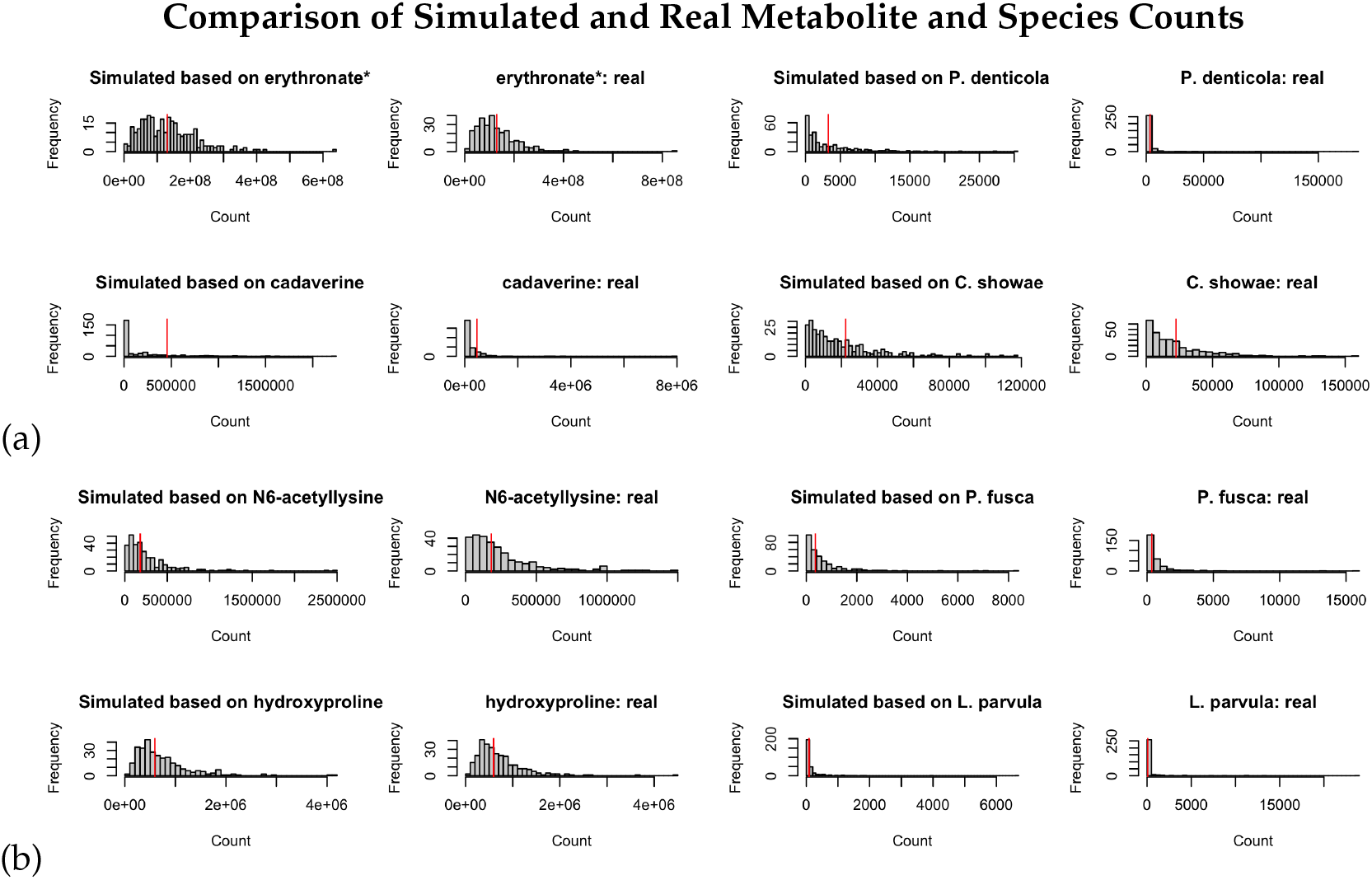
Evaluation of Goodness of Fit for the ZINB/NB and log-normal models by comparing the histogram of frequency between the simulated data and real data processed in Kraken2/Bracken. The 2nd and 4th columns are the real data with one species per figure. The 1st and 3 columns are the ZINB simulated data using the parameters estimated from the corresponding species. 1st and 2nd columns illustrate metabolites and 3rd and 4th columns present microbial species. a denotes ZINB/NB-based simulation and b denotes log-normal based simulation. (**a**) For two randomly selected metabolites and two randomly selected species (Kraken2/Bracken), comparison of simulated counts drawn from ZINB distribution (with parameters obtained from models fitted on the real data) and the real data. If the real data have less than 50 out of 289 zeros, the simulated counts are drawn from the negative binomial distribution with no zero inflation. Red vertical lines represent model-based means for each metabolite and species. (**b**) For two randomly selected metabolites and two randomly selected species (Kraken2/Bracken), comparison of simulated counts drawn from (ZI-)lognormal distribution (with parameters obtained from models fitted on the real data) and the real data. If the real data have no zeros, the simulated counts are drawn from the lognormal distribution with no zero inflation. Red vertical lines represent the log-scale means of the counts for each metabolite and species.

**Figure 3.**
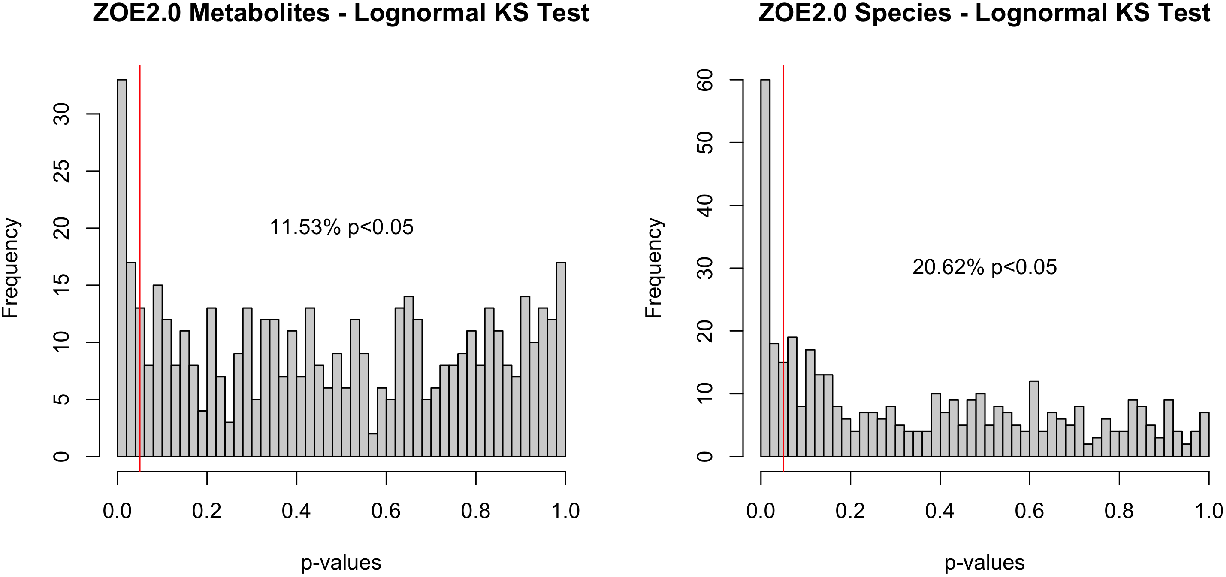
P-values obtained from lognormal (parameters from models fitted on nonzero counts for each metabolite and species) Kolmogorov-Smirnov test for ZOE 2.0 metabolites and Kraken2/Bracken microbiome species.

### 3.2. Estimation accuracy of underlying correlation in simulated correlated pairs of count data vectors

We evaluate the estimation accuracy of underlying correlations across our measures of correlation for each simulated pair of vectors. The four methods are (1) correlation based on the BZINB model (fitted with at most 1,000 E-M iterations); (2) correlation based on the BNB model (fitted with at most 1,000 E-M iterations); and (3) Pearson and (4) Spearman correlation on the vectors after elements were set to zero. For each of these simulations, the mean and median correlation approximations were based on 1,000 replicates.

In nearly all cases, BZINB and BNB-based correlations were closer to the true and theoretical correlation compared to the Spearman correlation (Figure 5 and Figure 6). As the number of zeros in either vector increased, the Spearman and model-based correlations tended to be lower than the true value. Similarly, as the theoretical correlation increased, the Spearman and model-based correlations also tended to be lower than the true value. These patterns were more noticeable with the Spearman correlation compared to the model-based correlations. BZINB-based correlations were more accurate than Spearman and BNB-based correlations in cases of high simulated underlying correlation or with more zeros, more noticeably when the simulated correlation was approximately 0.3 or higher.

**Figure 4.**
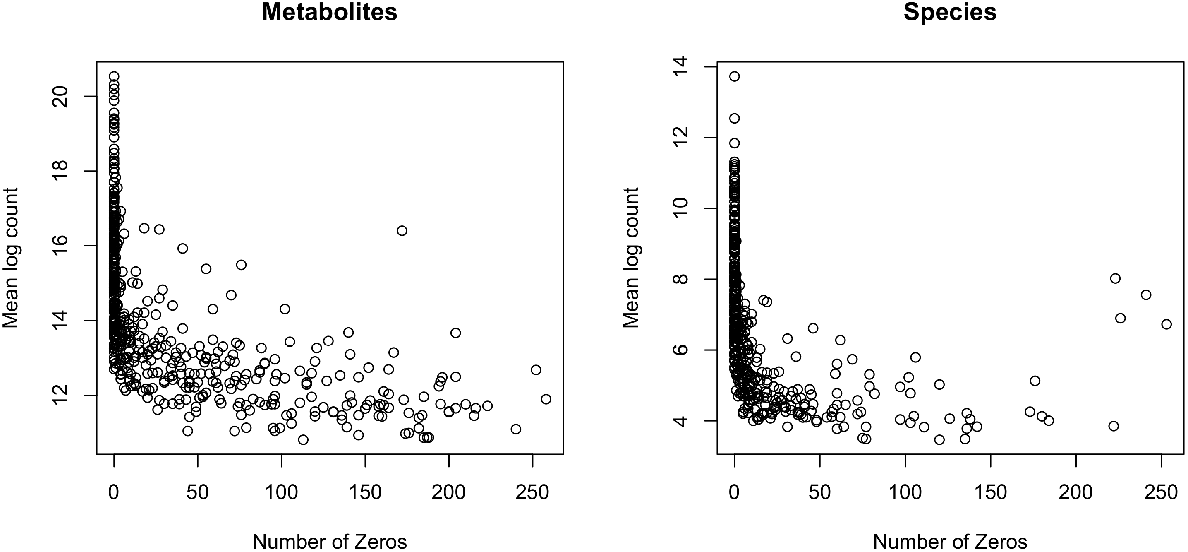
Number of zeros plotted against mean log nonzero count for each metabolite, and number of zeros plotted against mean log nonzero count for each Kraken2/Bracken species.

**Figure 5.**
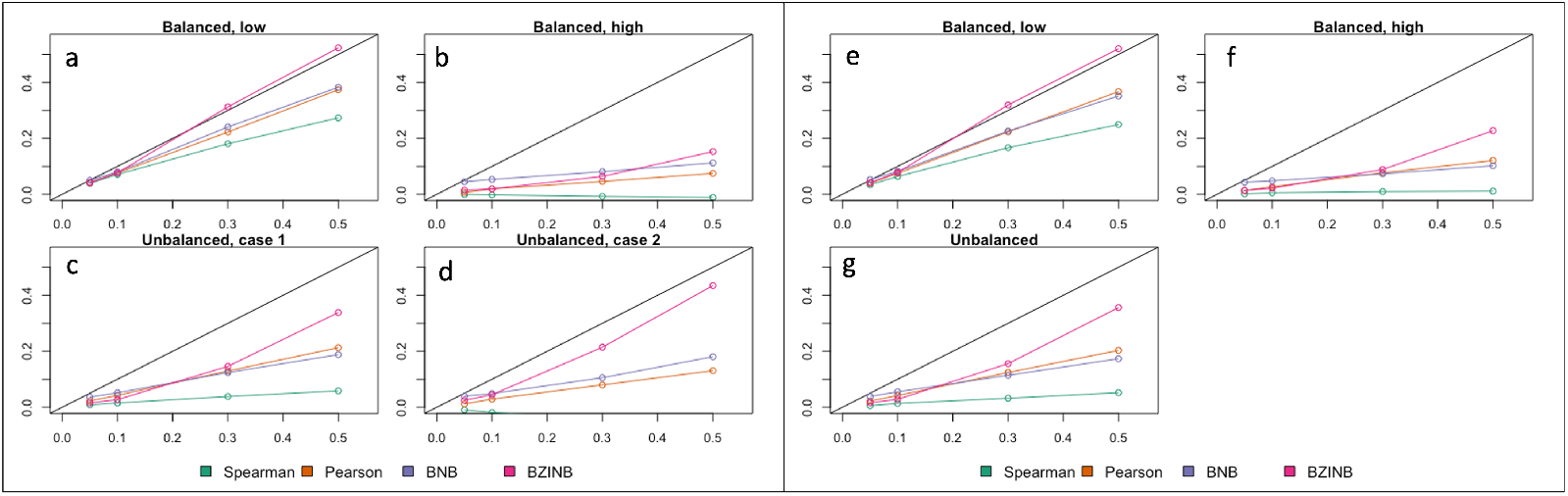
Mean approximated correlation for simulation of lognormal vectors representing pairs of metabolites and species corresponding to the (**a**) balanced, low, (**b**) balanced, high, (**c**) unbalanced, case 1, (**d**) unbalanced, case 2 expected numbers of zeros (parameters in Table 1; mean approximated correlation for simulation of lognormal vectors representing pairs of species corresponding to the (**e**) balanced, low, (**f**) balanced, high, (**g**) unbalanced expected numbers of zeros (parameters in Table 2). Each figure compares Spearman, Pearson, BNB-based, and BZINB-based correlations for five values of underlying correlation from the distributions where the simulated vectors are drawn from.

**Figure 6.**
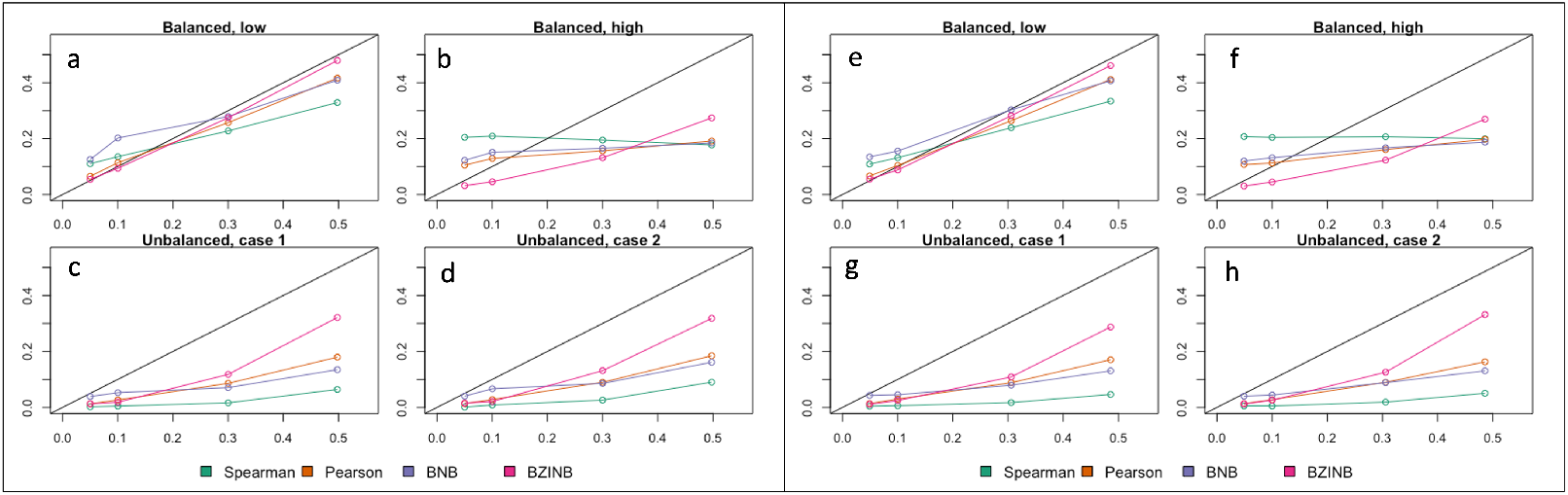
Mean approximated correlation for simulation of BZINB vectors representing pairs of metabolites and species corresponding to the (**a**) balanced, low, (**b**) balanced, high, (**c**) unbalanced, case 1, (**d**) unbalanced, case 2 expected numbers of zeros (parameters in Table 3 and Table A1); mean approximated correlation for simulation of BZINB vectors representing pairs of species corresponding to the (**e**) balanced, low, (**f**) balanced, high, (**g**) unbalanced, case 1, (**h**) unbalanced, case 2 expected numbers of zeros (parameters in Table 3 and Table A2). Each figure compares Spearman, Pearson, BNB-based, and BZINB-based correlations for five values of underlying correlation from the distributions where the simulated vectors are drawn from.

### 3.3. Accuracy evaluation of identified species modules using semi-parametric simulation

We sought to evaluate the accuracy of species modules identification using BZINB-based correlations compared to other correlations for spectral clustering. The ground truth was simulated using semi-parametric simulations as described in the Methods section. In the crafted semi-parametric simulated dataset representing counts for species belonging to 10 clusters (Figure 7a-b), we constructed affinity (distance) matrices using correlations from three methods (BZINB, BNB and Spearman correlations) in spectral clustering of species. To evaluate which method produces the most accurate and robust predicted 10 clusters, when different distance matrices were used, we compared (1) proportions of correctly predicted clusters, (2) the Adjusted Rand Index (ARI), and (3) the distance between the correlation matrices of the count matrices before and after adding cluster signals. For all resulting predicted clusters, there were instances when two or more separately assigned clusters were predicted to be essentially the same cluster (Figure 7c-e). This is likely due to the underlying similarities between species of different clusters in the original count data.

**Table 3.** lognormal simulations (metabolome-microbiome)

|  | Theoretical | Spearman | Pearson | BNB | BZINB |
| --- | --- | --- | --- | --- | --- |
| 1a | 0.5 | 0.273 (0.056) | 0.374 (0.115) | 0.383 (0.047) | 0.524 (0.058) |
| 1b |  | -0.011 (0.06) | 0.075 (0.122) | 0.112 (0.054) | 0.153 (0.21) |
| 1c |  | 0.058 (0.058) | 0.213 (0.118) | 0.188 (0.049) | 0.339 (0.18) |
| 1d |  | -0.064 (0.06) | 0.131 (0.123) | 0.181 (0.07) | 0.435 (0.205) |
| 2a | 0.3 | 0.181 (0.058) | 0.223 (0.096) | 0.241 (0.058) | 0.313 (0.08) |
| 2b |  | -0.007 (0.058) | 0.046 (0.099) | 0.081 (0.048) | 0.064 (0.127) |
| 2c |  | 0.038 (0.06) | 0.13 (0.101) | 0.124 (0.06) | 0.146 (0.144) |
| 2d |  | -0.043 (0.062) | 0.08 (0.093) | 0.106 (0.064) | 0.215 (0.171) |
| 3a | 0.1 | 0.071 (0.058) | 0.076 (0.071) | 0.08 (0.061) | 0.077 (0.068) |
| 3b |  | -0.002 (0.058) | 0.019 (0.071) | 0.053 (0.041) | 0.019 (0.05) |
| 3c |  | 0.015 (0.059) | 0.042 (0.071) | 0.052 (0.053) | 0.027 (0.053) |
| 3d |  | -0.019 (0.059) | 0.029 (0.07) | 0.049 (0.046) | 0.045 (0.07) |
| 4a | 0.05 | 0.039 (0.059) | 0.042 (0.065) | 0.05 (0.052) | 0.042 (0.048) |
| 4b |  | 0 (0.058) | 0.007 (0.063) | 0.045 (0.036) | 0.014 (0.042) |
| 4c |  | 0.009 (0.056) | 0.023 (0.061) | 0.036 (0.042) | 0.016 (0.034) |
| 4d |  | -0.01 (0.059) | 0.012 (0.065) | 0.04 (0.042) | 0.025 (0.045) |

**Table 4.** lognormal simulations (within microbiome)

|  | Theoretical | Spearman | Pearson | BNB | BZINB |
| --- | --- | --- | --- | --- | --- |
| 1a | 0.5 | 0.25 (0.058) | 0.367 (0.11) | 0.351 (0.047) | 0.52 (0.065) |
| 1b |  | 0.011 (0.061) | 0.121 (0.13) | 0.102 (0.046) | 0.228 (0.242) |
| 1c |  | 0.052 (0.06) | 0.203 (0.123) | 0.173 (0.051) | 0.356 (0.191) |
| 2a | 0.3 | 0.167 (0.06) | 0.223 (0.102) | 0.226 (0.056) | 0.319 (0.083) |
| 2b |  | 0.01 (0.057) | 0.077 (0.107) | 0.073 (0.041) | 0.089 (0.151) |
| 2c |  | 0.032 (0.059) | 0.124 (0.099) | 0.114 (0.054) | 0.156 (0.154) |
| 3a | 0.1 | 0.064 (0.059) | 0.075 (0.072) | 0.082 (0.061) | 0.078 (0.073) |
| 3b |  | 0.006 (0.06) | 0.027 (0.077) | 0.048 (0.037) | 0.022 (0.055) |
| 3c |  | 0.014 (0.057) | 0.042 (0.073) | 0.055 (0.046) | 0.028 (0.055) |
| 4a | 0.05 | 0.034 (0.059) | 0.04 (0.064) | 0.052 (0.051) | 0.041 (0.05) |
| 4b |  | 0.002 (0.061) | 0.013 (0.063) | 0.043 (0.035) | 0.014 (0.041) |
| 4c |  | 0.006 (0.059) | 0.022 (0.067) | 0.039 (0.04) | 0.016 (0.034) |

**Table 5.**
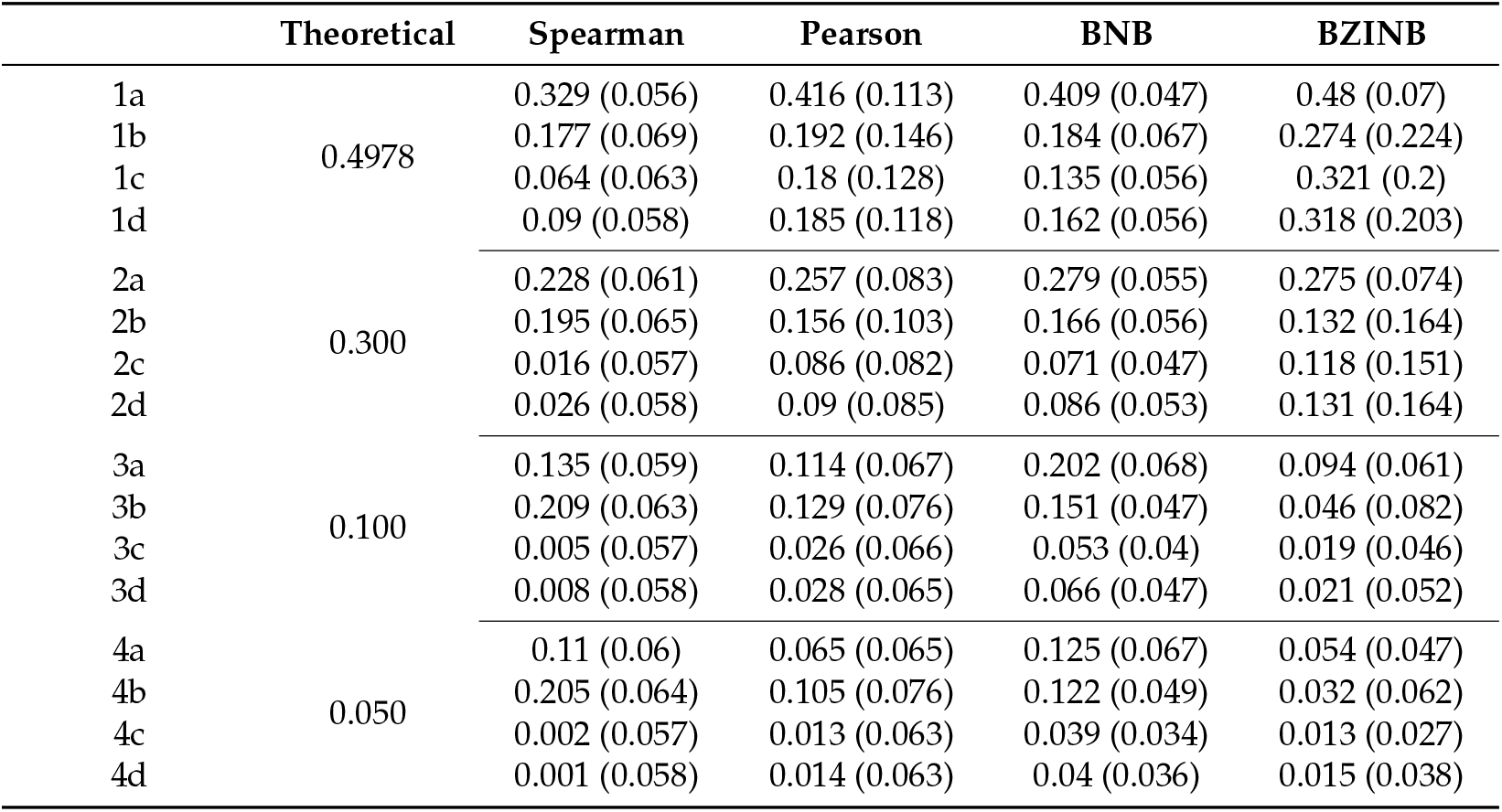
BZINB-based simulations (metabolome-microbiome)

**Table 6.**
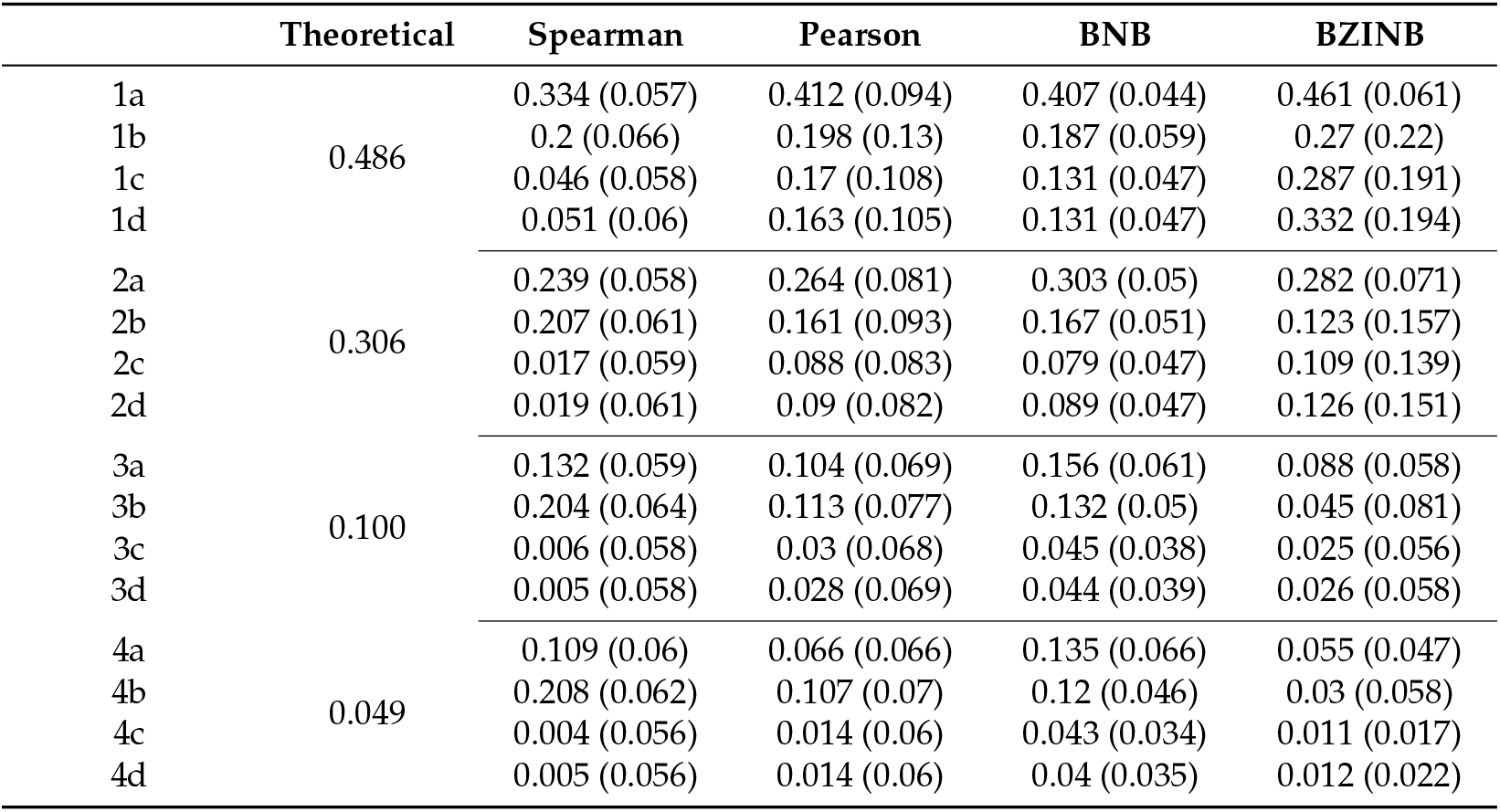
BZINB-based simulations (within microbiome)

|  | Theoretical | Spearman | Pearson | BNB | BZINB |
| --- | --- | --- | --- | --- | --- |
| 1a | 0.486 | 0.334 (0.057) | 0.412 (0.094) | 0.407 (0.044) | 0.461 (0.061) |
| 1b |  | 0.2 (0.066) | 0.198 (0.13) | 0.187 (0.059) | 0.27 (0.22) |
| 1c |  | 0.046 (0.058) | 0.17 (0.108) | 0.131 (0.047) | 0.287 (0.191) |
| 1d |  | 0.051 (0.06) | 0.163 (0.105) | 0.131 (0.047) | 0.332 (0.194) |
| 2a | 0.306 | 0.239 (0.058) | 0.264 (0.081) | 0.303 (0.05) | 0.282 (0.071) |
| 2b |  | 0.207 (0.061) | 0.161 (0.093) | 0.167 (0.051) | 0.123 (0.157) |
| 2c |  | 0.017 (0.059) | 0.088 (0.083) | 0.079 (0.047) | 0.109 (0.139) |
| 2d |  | 0.019 (0.061) | 0.09 (0.082) | 0.089 (0.047) | 0.126 (0.151) |
| 3a | 0.100 | 0.132 (0.059) | 0.104 (0.069) | 0.156 (0.061) | 0.088 (0.058) |
| 3b |  | 0.204 (0.064) | 0.113 (0.077) | 0.132 (0.05) | 0.045 (0.081) |
| 3c |  | 0.006 (0.058) | 0.03 (0.068) | 0.045 (0.038) | 0.025 (0.056) |
| 3d |  | 0.005 (0.058) | 0.028 (0.069) | 0.044 (0.039) | 0.026 (0.058) |
| 4a | 0.049 | 0.109 (0.06) | 0.066 (0.066) | 0.135 (0.066) | 0.055 (0.047) |
| 4b |  | 0.208 (0.062) | 0.107 (0.07) | 0.12 (0.046) | 0.03 (0.058) |
| 4c |  | 0.004 (0.056) | 0.014 (0.06) | 0.043 (0.034) | 0.011 (0.017) |
| 4d |  | 0.005 (0.056) | 0.014 (0.06) | 0.04 (0.035) | 0.012 (0.022) |

**Figure 7.**
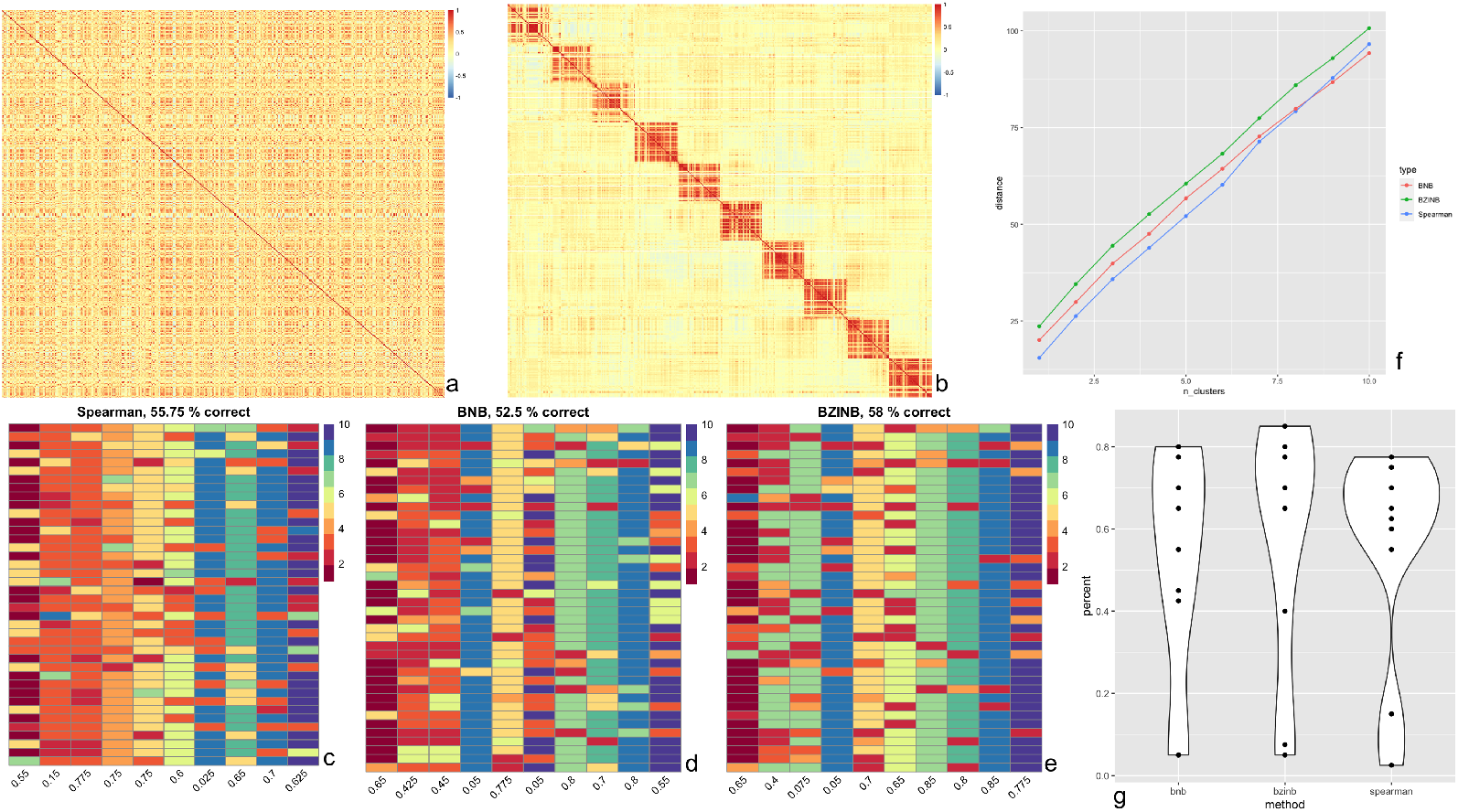
(**a**) Heatmap of BZINB-based correlation between Kraken2/Bracken counts of 400 of the species in ZOE2.0 in a random order; (**b**) Heatmap of BZINB-based correlation of the Kraken2/Bracken count data (in the same order as in (a)) after introducing simulated clusters; **(c-e**) Each column of cells represents a true cluster based on the simulation (b), and the colors represent the predicted clustering using affinity matrix made from (c) BNB, (d) BZINB, and (e) Spearman correlations; (**f**) Distance (Frobenius norm) between the correlation matrices of nested predicted clusters between data with (as in Figure 7b) and without (as in Figure 7a) increased correlations that represented the clusters: the first set (number of clusters = 1) is the predicted cluster with the greatest distance between correlation matrices. For each increase in the number of clusters, we included an additional cluster in the order of decreasing distances. This was done using the BNB-based, BZINBbased, and Spearman correlation matrices and their corresponding cluster predictions; (**g**)Violin plot of cluster-wise percent accuracy for each of the 10 clusters, comparing BNB, BZINB, and Spearman correlation-based affinity matrices.

Firs, while several approaches exist to quantify clustering accuracy, we considered the proportions of species in each assigned cluster that were predicted to be in the same cluster. We found that in the data with simulated clusters (simulated as in Methods Section 2.4.2), using the BZINB-based correlation resulted in the highest overall proportion of accurate cluster assignments, while the BNB-based correlation resulted in the lowest accuracy (Figure 7g). Clusters that were generated using BZINB correlations had up to 85% accuracy, and most had at least 65% accuracy. On the other hand, most of the Spearman correlation-based clusters had between 55% to 75% accuracy. There was a moderate percentage (40-55%) of inaccurately predicted BNB correlation-based clusters.

Second, we evaluated the accuracy of the predicted clusters for each correlation type using the ARI. Higher ARI indicates higher consistency between the observed and the simulated cluster membership. In concordance with the proportion of accurate cluster assignments, the affinity matrix based on the BZINB-based correlation resulted in an ARI of 0.43, which was the highest among the three. The ARI for the BNB-based and Spearman correlations were 0.38 and 0.34 respectively. Therefore, BZINB model-based clustering provides the best clustering results.

Third, we compared the three methods according to the distance between correlation matrices. The distance between two correlation matrices (where BZINB correlations were calculated for each pair of species) with partitions representing clusters is one way to compare networks of microbial species or other multi-omics between two health/disease groups. Further, distances between correlation matrices of two health/disease states within each species-cluster allows the determination of clusters that are differentially inter-correlated between these conditions.

Different types of correlation measurements vary in terms of power for detecting between-network differences. Therefore, to compare the correlation types in quantifying the difference between a network with clusters of highly correlated species and a network with clusters of weakly correlated species, we computed distances between the two networks for nested sets of clusters. The first set was the cluster with the greatest distance, and we proceeded by sequentially adding clusters in order of decreasing distances. We used the Frobenius norm of the absolute difference between the correlation (sub-)matrices as the distance measure because it accounts for all matrix entries and is easily understood as an extension of the Euclidean distance between vectors. This was done using the BNB-based, BZINB-based, and Spearman correlation matrices and their corresponding cluster predictions. Distances between two correlation networks were consistently maximized using BZINB correlations, while they were the lowest using Spearman correlations for all but one of the cluster sets (Figure 7f).

### 3.4. Application in in the ZOE2.0 study

#### 3.4.1. Interactions among commensal species and among ECC-associated species

The most abundant species in a microbial community are of natural interest when examining microbial community dynamics in dysbiotic conditions such as those leading to the development of dental caries development. They represent a group of commensal species that may be perturbed in the presence of dental disease. Between the top 10 most abundant species in ZOE2.0, there are stronger correlations in the context of disease (ECC group) compared to the caries-free (non-ECC) group (Figure 8). The Spearman, BNB, and BZINB-based correlations between the 10 most abundant species are very similar because these species have no missing counts. In contrast, when one or more species have higher proportions of zeros, there may be a larger difference between the BNB and BZINB correlations. This is in accordance with simulation results, where all the correlation types were similar under few zeros in both vectors, while the different correlation types were less similar when there were excess zeros in one or more of the vectors

**Figure 8.**
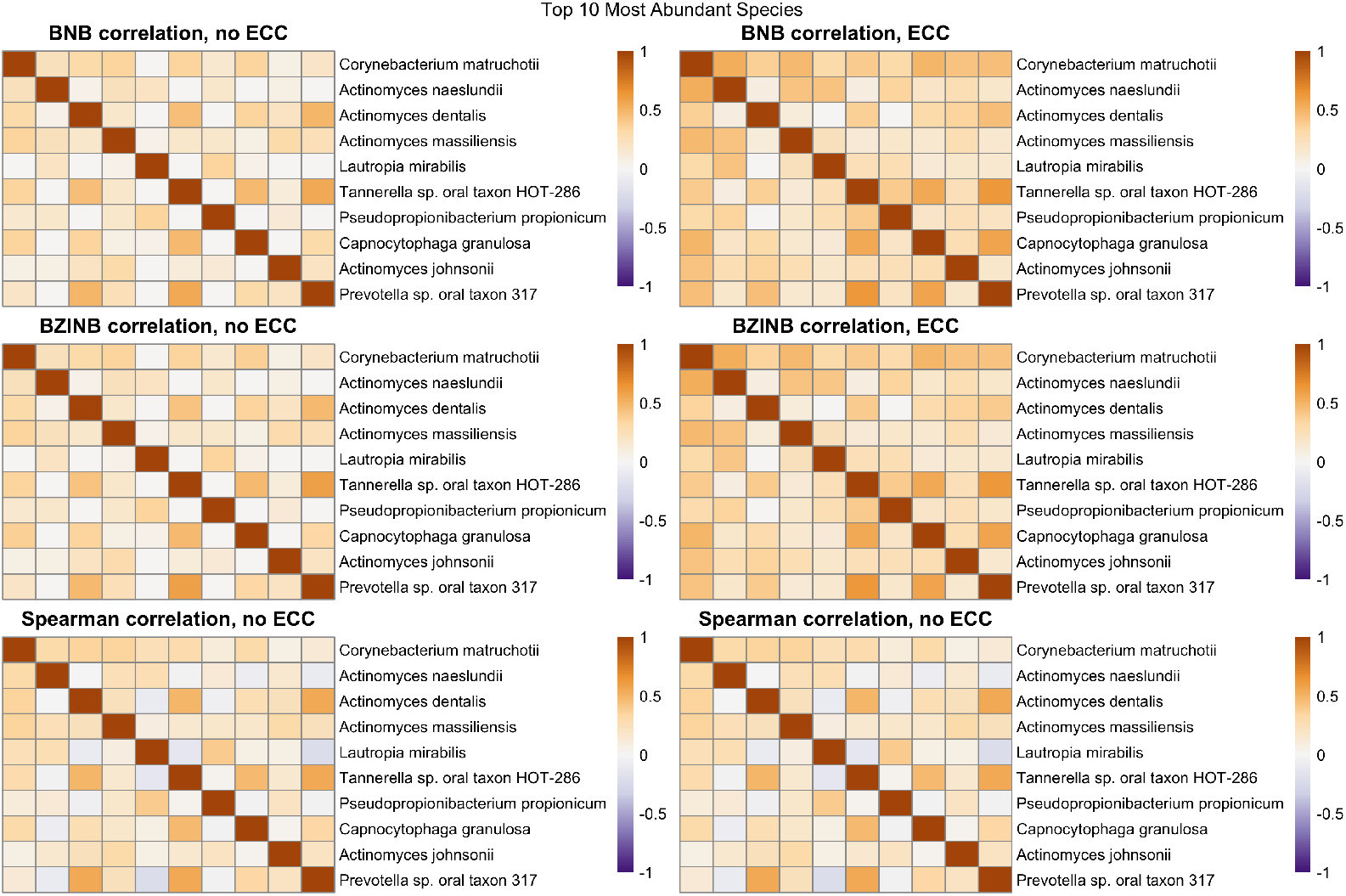
Heatmaps of BZINB-based and Spearman correlations between the top 10 species with the highest overall abundance for each health/dental disease group (non-ECC versus ECC) in the ZOE2.0 Kraken2/Bracken data.

We also examined interactions between metabolites and species that have been previously shown to be strongly associated with the presence of ECC. Therefore, next, we focused on the set of 15 metabolites and 16 species that have been previously identified to be associated with ECC in differential abundance analyses [5,27]. To understand these ECC-associated interaction networks/pathways, we compared correlations of between-species networks and species-metabolite networks as follows. First, we compared BZINB-based (Figure 10) and Spearman-based correlation between-species networks (Figure 11). We found that *Veillonella atypica* is highly correlated with several ECC-associated *Prevotella* species among children affected with ECC using both of these correlations (Figure 10 and Figure 11).On the other hand, many of these *Prevotella* species tend to be strongly correlated with *Leptotrichia, Lachnospiraceae*, and *Lachnoanaerobaculum* species in children unaffacted by ECC. This points to two possible co-abundance patterns: one where *Prevotella, Leptotrichia, Lachnospiraceae*, and *Lachnoanaerobaculum* taxa may coexist in biofilms without disease, and another pattern of mutual benefit among *V. atypica* and *Prevotella* species when disease is present. In this case, the co-abundance pattern between these two species can be explained by their beneficial interrelation in metabolic activities: carbohydrates and sugar alcohols from the diet are subjected to glycolysis, which creates anaerobic conditions by consuming oxygen, and produces pyruvate that can be converted into lactate by *Prevotella* species. On the other hand, *Veillonella atypica* is an anaerobic bacteria that uses lactate as their sole carbon source, converting into weaker acids, such as acetate and propionate [32]. Between the 15 species of interest, the BZINB correlation network included only one strong correlation involving *Streptococcus mutans* and *Veillonella atypica* in healthy subjects, whereas the Spearman correlation network did not include *Streptococcus mutans* at all. *Streptococcus* and *Veillonella* species are very common in the supragingival oral biofilm, and Mashima *et al*. 2015 showed a *Streptococcus*-*Veillonella* link in early dental plaque formation–in fact, *Streptococcus mutans* is well-known as a major lactic acid producer from the fermentation of dietary carbohydrates, which benefits *Veillonella* species since it utilize lactate produced by *Streptococcus mutans* and converts it into weaker acids, such as acetate and propionate contributing to acid neutralization. Therefore, the identified strong correlation between the two species is reasonable. However, when acid production occurs at a greater rate and frequency than that of acid neutralization, dental caries will develop. So, in subjects with caries, *Veillonella atypica* was more abundant compared to those without caries (Figure 9). Therefore, the *Streptococcus mutans*-*Veillonella atypica* dynamic may be somewhat overpowered by *Streptococcus mutans* once ECC develops.

**Figure 9.**
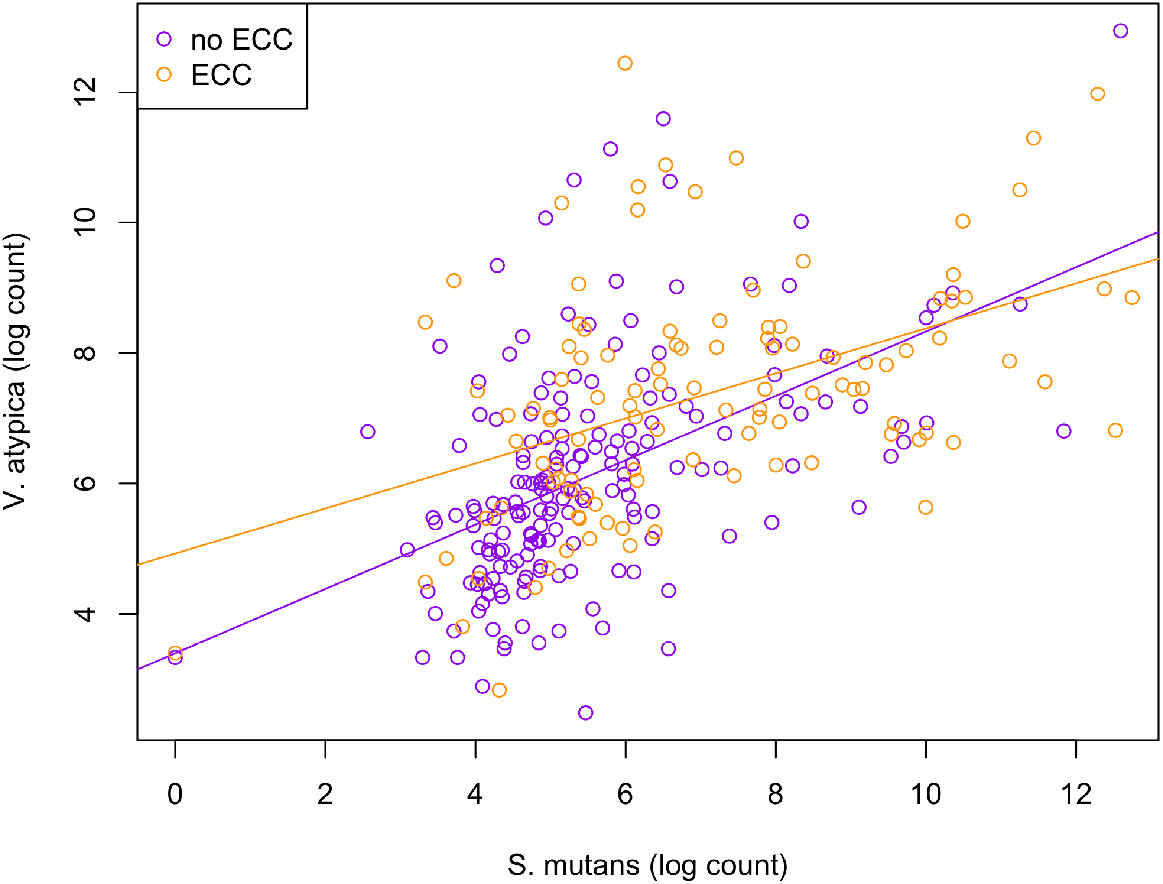
Scatterplot illustrating the comparison of relationships between S. mutans and V. atypica abundances between health (non-ECC) and disease (ECC) groups.

**Figure 10.**
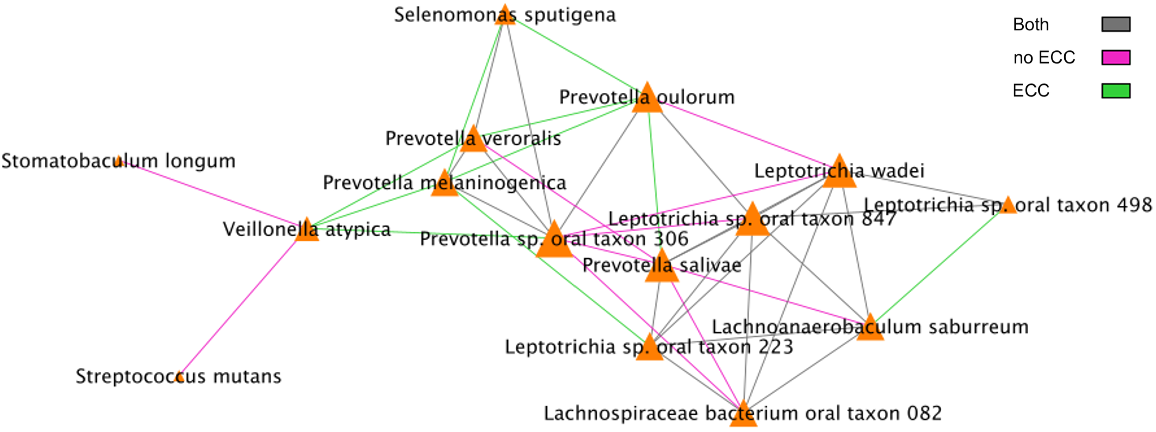
BZINB correlations between species. The strongest 30% of correlations are included in the diagrams, and the color of the lines represent whether the correlation was strong in one or both of the health/disease groups.

**Figure 11.**
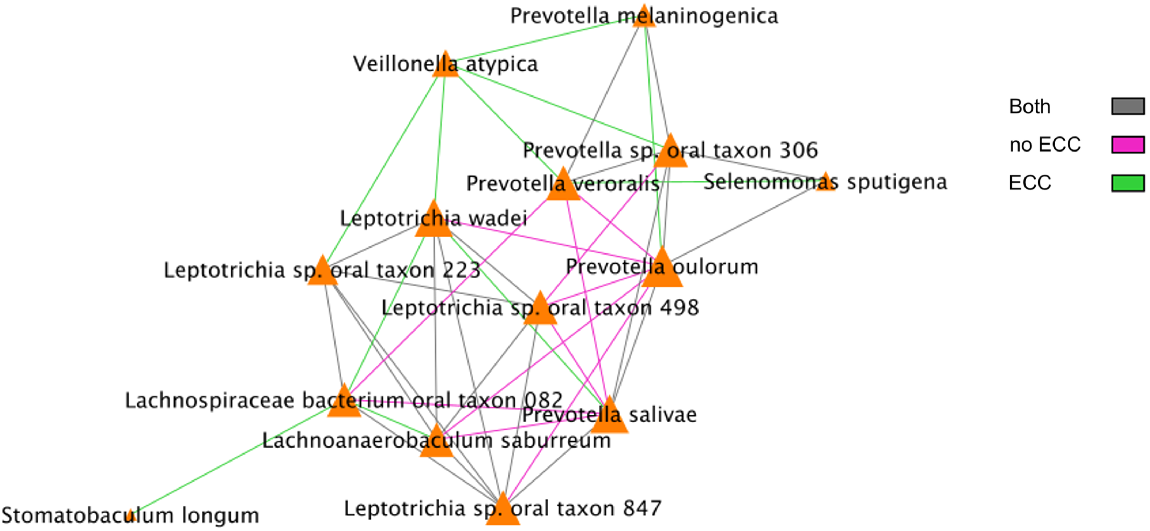
Spearman correlations between species. The strongest 30% of correlations are included in the diagrams, and the color of the lines represent whether the correlation was strong in one or both of the health/disease groups.

Additionally, we compared BZINB-based (Figure 12) and Spearman correlations-based species-metabolite networks (Figure 13). In the oral biofilm, when diet-associated carbohydrates are present, carbohydrate-degrading species tend to increase in abundance and the local environment pH may decrease [29]. To observe the differences in species that are highly correlated with carbohydrates of interest in healthy subjects and subjects with ECC, we focused on interpretation of four carbohydrates that were previously shown to be significantly and positive associated with ECC in Heimisdottir *et al*. 2021. We used the BZINB-based correlations because some of the species had excessive zeros. For each of the five carbohydrates, we compared the strongest 5% of metabolite-species correlations between health/disease groups. In caries-affected participants, the amount of three of the carbohydrates (fucose, sedoheptulose-7-phosphate, and N-acetylneuraminate) is strongly correlated with many *Prevotella* species. According to Takahashi *et al*. 2005, *Prevotella* neutralizes pH but may also favor the presence of other pathogenic species. In healthy subjects, we found the carbohydrates to be correlated with *Streptococcus, Fusobacterium*, and *Selenomonas* species, many of which have been described as carbohydrate-degrading or pH-neutralizing in the oral biofilm [30,31], or are a core part of the normal flora. In the BZINB network, 3-(4-hydroxyphenyl)lactate (HPLA) had many strong correlations with various species in participants with ECC but much less among unaffected ones. HPLA is a metabolite in the tyrosine metabolism pathway that functions similarly to lactate, which has been previously shown to be an important metabolic regulator in multiple pathways (including glucose metabolism) in various parts of the human body [34,35]. The differing strengths of correlations in the two health/disease groups could indicate that HPLA is metabolized differently by ECC-associated species in the context of a dental caries-promoting environment, and may be a candidate for further investigation in its role in ECC development. Furthermore, HPLA is strongly associated with many *Streptococcus* species in healthy subjects and with many *Prevotella* species among those with ECC, similarly to what was found for ECC-associated carbohydrates.

**Figure 12.**
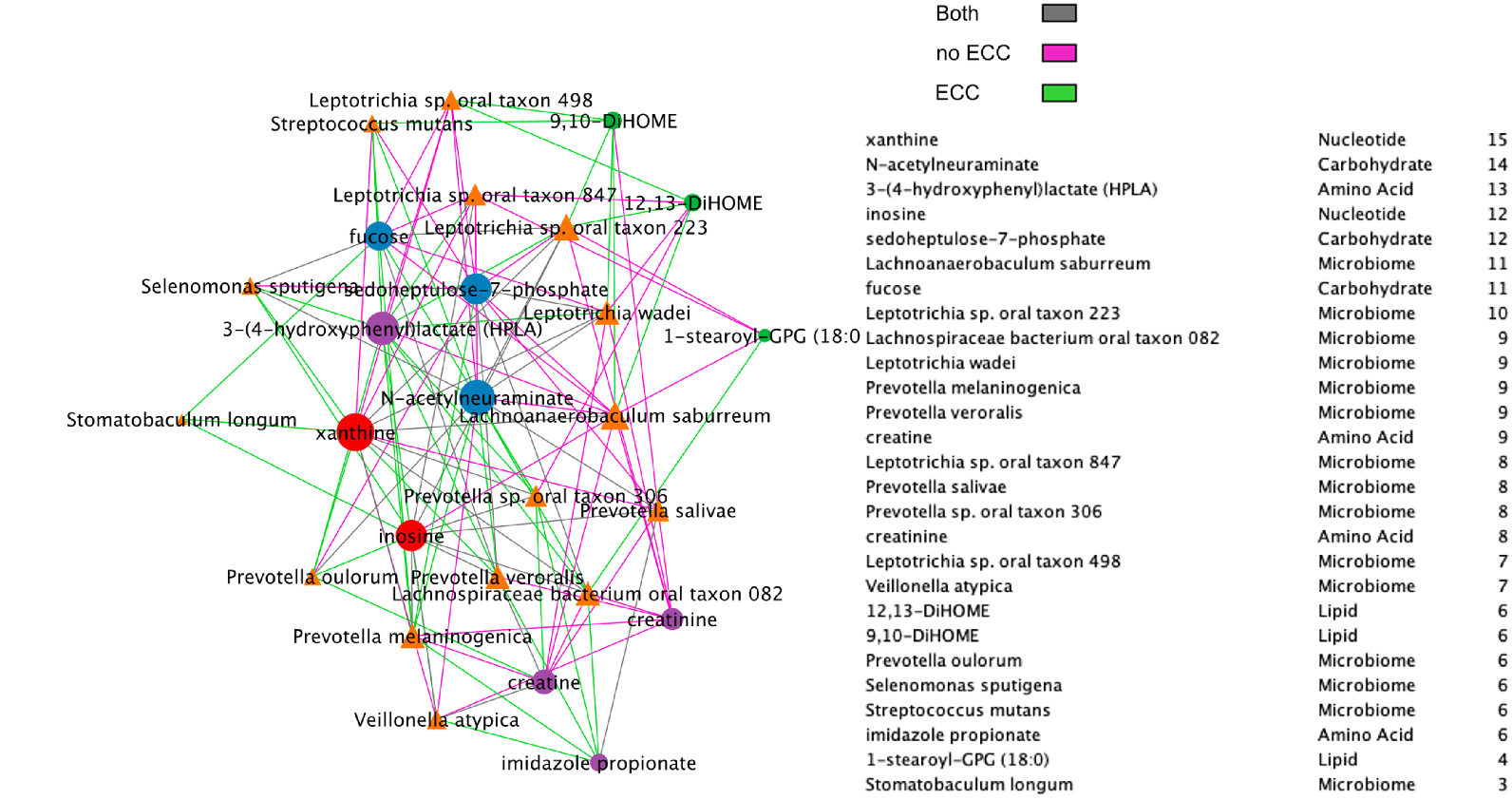
BZINB network between species and metabolites including a node degree table. The strongest 30% of correlations are presented in the diagrams and line colors represent whether the correlation was strong in one or both of the health/disease groups.

**Figure 13.**
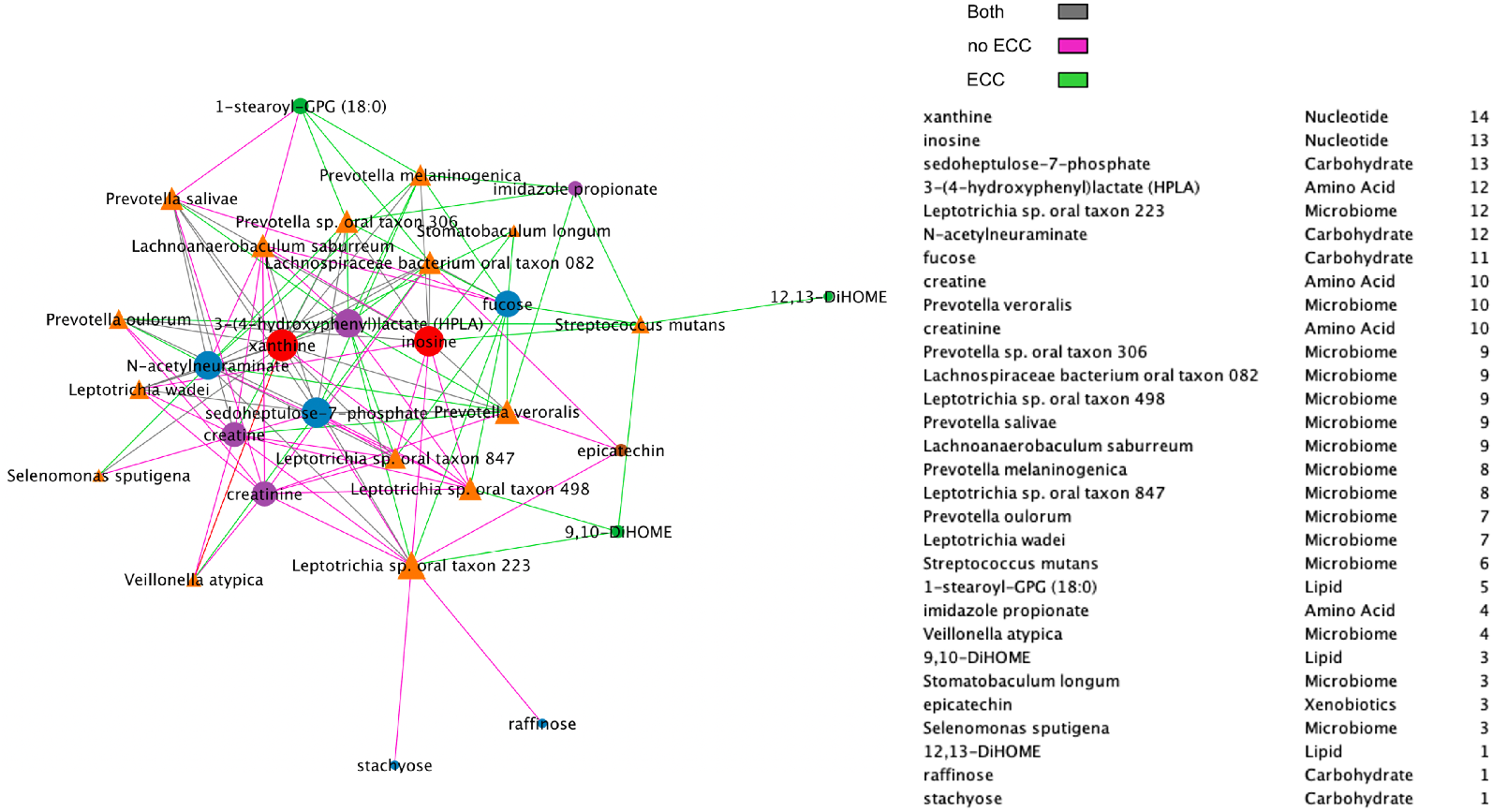
Spearman network between species and metabolites, presenting positive correlations only and including a node degree table. The strongest 30% of positive correlations are presented in the diagrams, and line colors represent whether the absolute correlation was strong in one or both of the health/disease groups.

Overall, Spearman and Pearson correlations are not suitable for data with excess zeros because Spearman is influenced by ties and Pearson requires a linear association. The negative binomial distribution accounts for the presence of zeros, which makes the BNB distribution a better choice for modeling the relationship between a typical pair of species or metabolites. When there are excess zeros in either or both species or metabolites in a pair, the BZINB model can account for the zero-inflation while approximating the correlation of the nonzero components.

### 3.5. Species modules identified using BZINB-based correlation and spectral clustering

We applied cut-based spectral clustering to the ZOE 2.0 data separately for each health/disease group. We compared results between BZINB-based and Spearman correlations when constructing the affinity matrix. To determine the optimal number of clusters, we plotted the eigenvalues of the graph Laplacian for each affinity matrix (Appendix Figure 5). According to the eigengap method [26], the optimal number of clusters was 2 for each affinity matrix; for more interpretable results, we set the number of clusters to be 6 in each case. To visualize the results of cut-based clustering, we created heatmaps of standardized counts for all species, where the species are grouped and annotated by predicted cluster and the study participants are annotated according to health/disease and batch group. There were visible within-cluster similarities and differences between the clusters for count patterns (Figure 14). Many species that were predicted to be in the second and fifth (shown in blue and orange, respectively, in the top bars of Figure 14) in the healthy group had been classified in third cluster (shown in green) in the disease group. In other words, some species that were more similar to the first and fifth clusters in the healthy group were instead more similar to the third cluster in subjects with ECC. The different co-varying patterns in these species that may be a reflection of differences in the microbial community structure and function in ECC.

**Figure 14.**
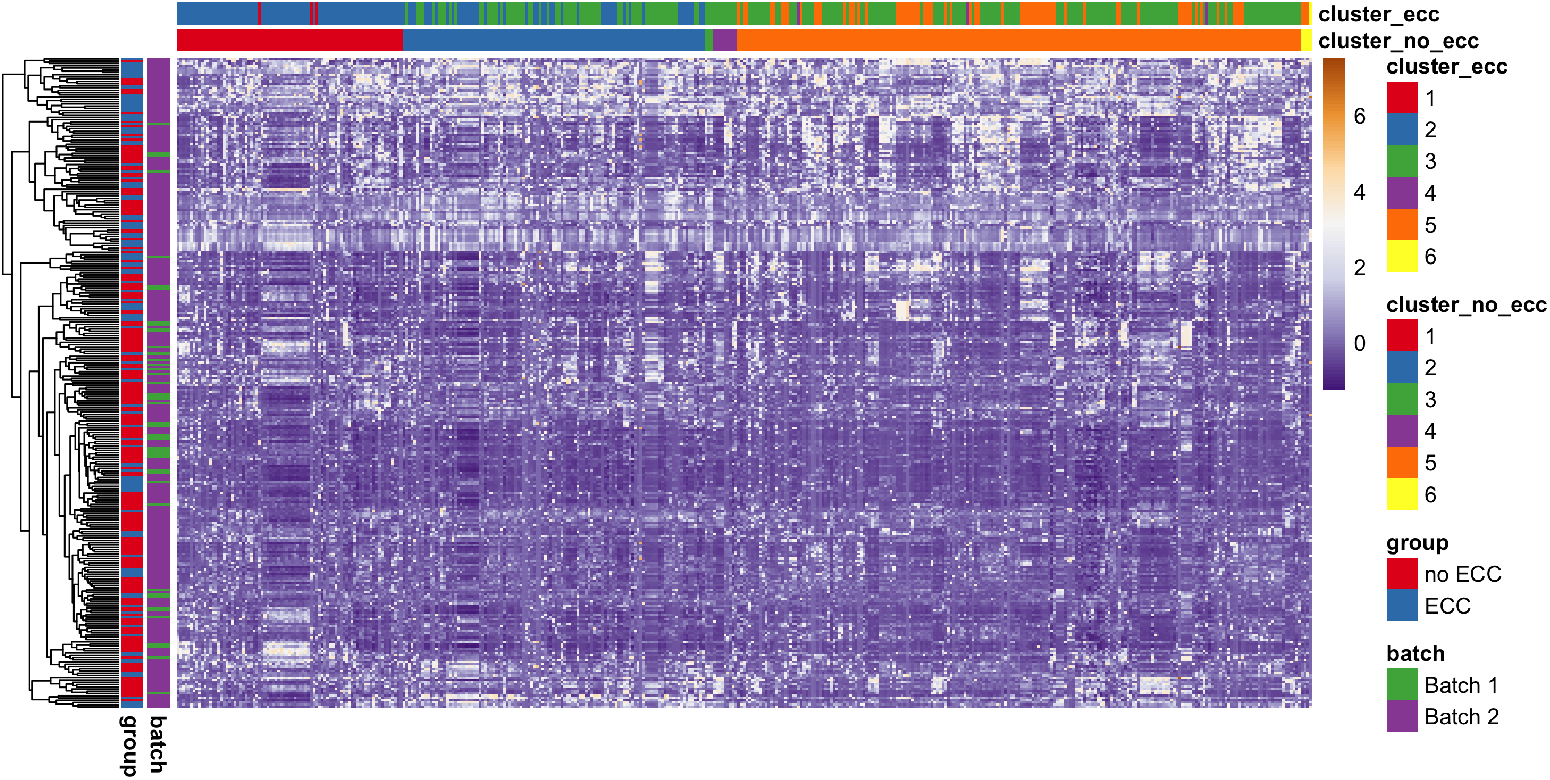
Heatmap of species abundance illustrating species module identification results (species are columns, modules are presented with different colors) using BZINB-based species spectral clustering. Each column represents a single species. Columns are ordered by the clusters predicted from the affinity matrix based on the BZINB correlations between species in the healthy (non-ECC) group. The columns are annotated to show and compare the estimated clusters within health (non-ECC) and disease (ECC) groups. Each row represents a participant, and the rows are ordered based on hierarchical clustering. Rows (n=289) are annotated to denote health/disease and batch groups. Standardized counts were used.

## 4. Discussion

In this paper, we introduce a new method BZINB-iMMPath entailing a bivariate zero-inflated negative binomial (BZINB) model-based correlation for network analysis of pairs of vectors of omics count data and module identification. The model makes reasonable assumptions regarding dropouts and excess zeros as structural zeros in the observed microbiome data compared to other types of zeros. Therefore, the microbial correlation distribution is assumed to be that of the latent bivariate negative binomial model. Our approach improves the estimation of correlations compared to the traditional Pearson correlation and the more robust Spearman’s rank correlation coefficients. In contrast to Pearson and Spearman correlations, the BZINB model accommodates zeros in a flexible manner (in either or both vectors of each pair) and estimates the correlation under the bivariate negative binomial model. For each pair of omics features, the BZINB model is fitted and a model-based correlation is computed from the estimated parameters. Using the model, we can calculate the correlations between pairs of omics features in the same layer (i.e., between pairs of microbial species) or between two different layers (i.e., between pairs of metabolites and species). These correlations may then be used in other applications such as networks’ visual representations and identification of clusters of omics features. Accordingly, we applied the new method to microbial species and metabolite data obtained in an oral microbiome study of early childhood dental disease. Using visual comparisons and goodness-of-fit tests, we determined that the negative binomial and lognormal distributions were appropriate for modeling most metabolites and species. In addition to accounting for zero inflation, marginally, the negative binomial distribution is a natural choice to model count data. Therefore, our model-based correlation approach has several advantages over conventional measures of correlation when applied to bivariate count data with excess zeros. In addition, correlations estimated from BZINB can be used as the affinity matrix in the cut-based spectral clustering method for species module identification in zero-inflated microbiome data. Modules can be compared between groups of interest (e.g., health versus disease) and help identify species that demonstrate important between-group pattern differences.

To evaluate the performance of BZINB-iMMPath, we used real data-inspired simulations to estimate the accuracy of underlying correlations in microbiome data; real data-based semi-parametric simulations to access the accuracy of module identification; and finally, we applied it in a sizeable oral microbiome study to identify ECC-associated microbial networks and modules. Specifically, we simulated pairs of count vectors representing typical metabolite and microbial species vectors from ZOE 2.0 to compare the accuracy of Spearman, BNB, and BZINB model-based correlations. We fitted the BZINB model to each metabolite-species and species-species pair to construct visualizations of ECC disease group-specific filtered networks and build affinity matrices for cut-based spectral clustering. Using the simulated vector pairs, the BZINB model-based correlation was on average closer to the underlying correlation when there were more zeros in one or both vectors compared to the Spearman correlation coefficient. Notably, the average BZINB-based correlation was higher than the other correlation types when the underlying correlation was high (>0.3) and when there was zero inflation in at least one of the vectors. Therefore, we recommend using the BZINB-based correlation for the identification of strongly correlated pairs when zero inflation is present. The application in ZOE 2.0 not only highlighted previously known networks involving carbohydrate metabolites but also revealed novel regulation relationships between species and metabolites, and ECC-associated species modules.

The most noticeable limitation of the new approach is that the BZINB model allows for only positive model-based correlations. Ideally, the off-diagonal entries in the covariance matrix in BZINB should allow both positive and negative values. However, in most omics contexts, positively correlated features are arguably of most of interest. For example, in gene expression data, the vast majority of genes do have positive or near-zero correlation [36,37]. Positive correlations among bacterial species are also more common compared to negative correlations (Figure 1). Of course, there are cases where negative correlations are of interest, for example in the context of species competition, other correlation measures could be used. However, incorporating negative correlations can introduce another layer of complexity to network analysis applications for multi-omics and cluster identification. For example, negative correlations may be considered with different importance compared to positive correlations. Further, negative correlations within one layer of omics (such as microbiome), which could represent competition, may be more of interest compared to negative correlations between layers (e.g., microbiome and metabolome), which could be more complex in terms of direction of influence. This leaves room for future methods development, for example, wherein other bivariate (or multivariate) models can be evaluated in terms of goodness-of-fit for certain types of omics data that could accommodate negative correlations. Meanwhile, identifying positive correlations between bacteria and metabolites is a logical priority, because of biological interest regarding 1) which bacteria generate or up-regulate which metabolites, and 2) which biochemicals are associated with bacterial abundance (e.g., possibly growth). Meanwhile, negative correlation (like inhibition or competition) is harder to interpret as detailed above, and in our BZINB model, positive correlations are presented as such and negative correlations are estimated as near-zero.

In our application to the ZOE 2.0 study microbiome data, we determined that (1) there were relatively fewer zero counts when taxa were identified through the ora health-specific Kraken2/Bracken pipeline, compared to the data from the still widely used HUMAnN 2.0 pipeline; (2) zero inflation does not appear to be a significant issue for many of the named metabolites; and (3) in the absence of excess zeros, other measures of correlation appear to be just as adequate as the BZINB-based correlation. Because HUMAnN 2.0 generated data are very sparse, our method is even more powerful in those data, as well as similarly sparse gene-level metagenomics or metatranscriptomics data.

In sum, in this paper we demonstrate that the new method based on the BZINB model is a useful alternative to Spearman or Pearson correlation in estimating underlying correlations for bivariate count data that are zero-inflated in one or both dimensions. Because the model accommodates both technical and true zeros, it is suitable for multiomics data types including microbiome and metabolome. To identify differences between health/disease groups, we prioritized and illustrated the strongest correlations within each group, allowing the visualization of important dynamic relationships and their between-group comparison. Finally, these correlations can also be used in identifying modules, i.e., clusters of correlated metabolites and microbial species, which could be of biological interest both in terms of disease pathogenesis and intervention targeting.

## Author Contributions

Conceptualization, B.L. and D.W.; methodology, B.L, H.C. and D.W.; software, B.L, H.C. and C.L.; validation, B.L. and A.A.R.; formal analysis, B.L.; resources, K.D., J.R. and C.L.; data curation, K.D.; writing—original draft preparation, B.L and D.W.; writing—review and editing, B.L., H.C, K.D., A.A.R. and D.W.; visualization, B.L. and D.W.; supervision, D.W. All authors read and agreed to the published version of the manuscript.

## Institutional Review Board Statement

The study was conducted in accordance with the Declaration of Helsinki, and approved by the Institutional Review Board (or Ethics Committee) of University of North Carolina-Chapel Hill (14-1992, latest approved on 21 February 2022).

## Informed Consent Statement

Written informed consent was obtained from legal guardians of all children who participated in the ZOE 2.0 study.

## Data Availability Statement

ZOE 2.0 microbiome data are publicly available in the dbGaP repository at https://www.ncbi.nlm.nih.gov/gap under the umbrella study name Trans-Omics for Precision Dentistry and Early Childhood Caries or TOPDECC (accession: phs002232.v1.p1) via the Sequence Read Archive (SRA) Bioproject PRJNA671299 at https://www.ncbi.nlm.nih.gov/bioproject/671299. Metabolomics raw spectral data have been made publicly available via the MetaboLights repository project MTBLS2215 at https://www.ebi.ac.uk/metabolights/MTBLS2215. The code used for analysis is available at https://github.com/blin24/BZINB-iMMPath.

## Acknowledgments

The authors would like to thank ZOE 2.0 study participants for their contributions.

## Abbreviations

The: following abbreviations are used in this manuscript:
ZINB: Zero inflated negative binomial
BNB: Bivariate negative binomial
BZINB: Bivariate zero inflated negative binomial
MI: Mutual Information
ECC: Early childhood caries

## Appendix A

**Table A1.**
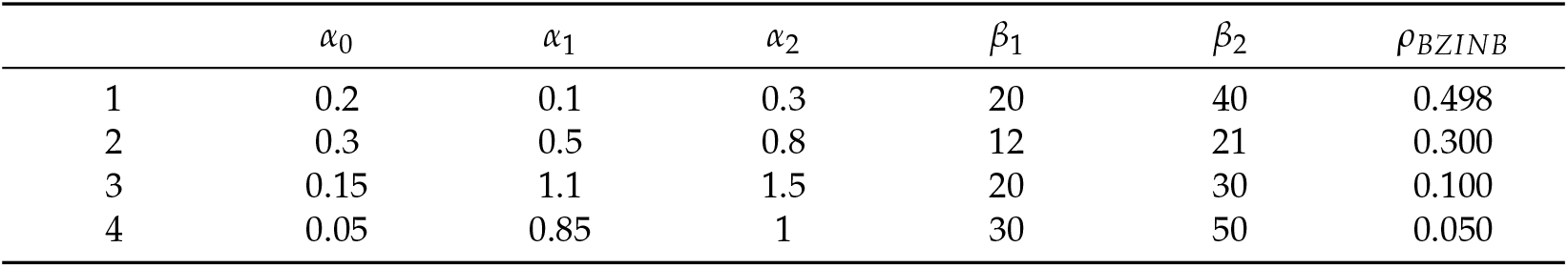
Shape and scale parameters used to obtain various values of correlation for BZINB simulation for vectors pairs that represent pairs of metabolites and species

| | $\alpha_0$ | $\alpha_1$ | $\alpha_2$ | $\beta_1$ | $\beta_2$ | $\rho_{BZINB}$ |
| --- | --- | --- | --- | --- | --- | --- |
| 1 | 0.2 | 0.1 | 0.3 | 20 | 40 | 0.498 |
| 2 | 0.3 | 0.5 | 0.8 | 12 | 21 | 0.300 |
| 3 | 0.15 | 1.1 | 1.5 | 20 | 30 | 0.100 |
| 4 | 0.05 | 0.85 | 1 | 30 | 50 | 0.050 |

**Table A2.** Shape and scale parameters used to obtain various values of correlation for BZINB simulation for vectors pairs that represent pairs of species

| | $\alpha_0$ | $\alpha_1$ | $\alpha_2$ | $\beta_1$ | $\beta_2$ | $\rho_{BZINB}$ |
| --- | --- | --- | --- | --- | --- | --- |
| 1 | 0.3 | 0.3 | 0.3 | 30 | 40 | 0.486 |
| 2 | 0.35 | 0.7 | 0.8 | 20 | 30 | 0.306 |
| 3 | 0.1 | 0.75 | 1 | 30 | 30 | 0.100 |
| 4 | 0.05 | 0.9 | 1 | 50 | 50 | 0.049 |

**Figure A1.**
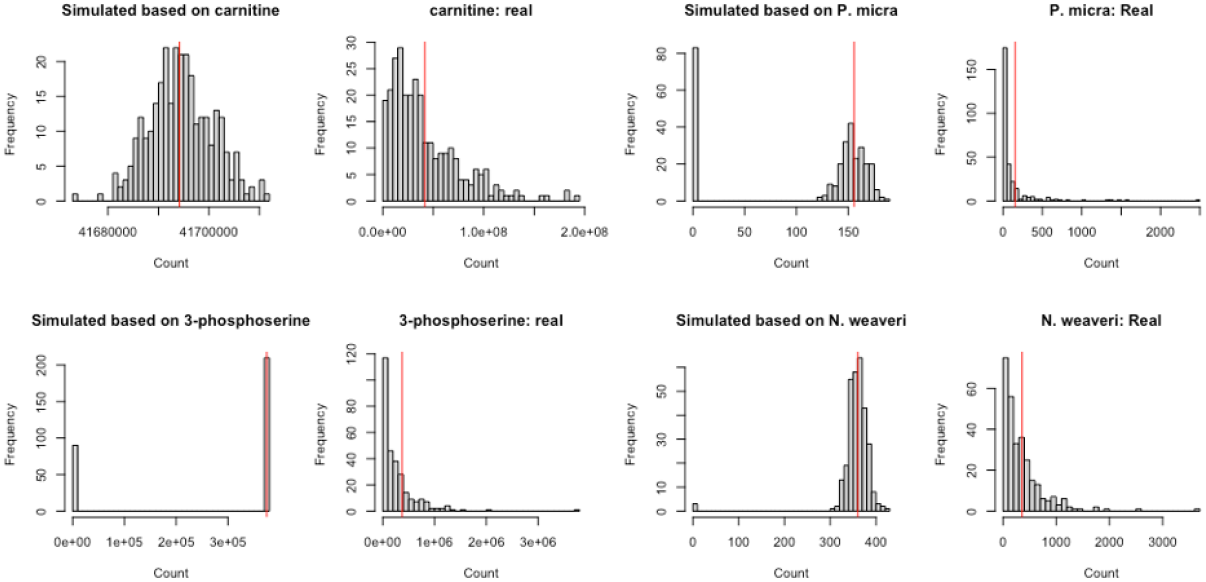
Comparison of simulated counts drawn from (ZI-)Poisson distribution (with parameters from model fitted on the real data) and real data of 4 randomly selected metabolites and species. Red vertical lines represent the model-based means for each metabolite and species.

**Figure A2.**
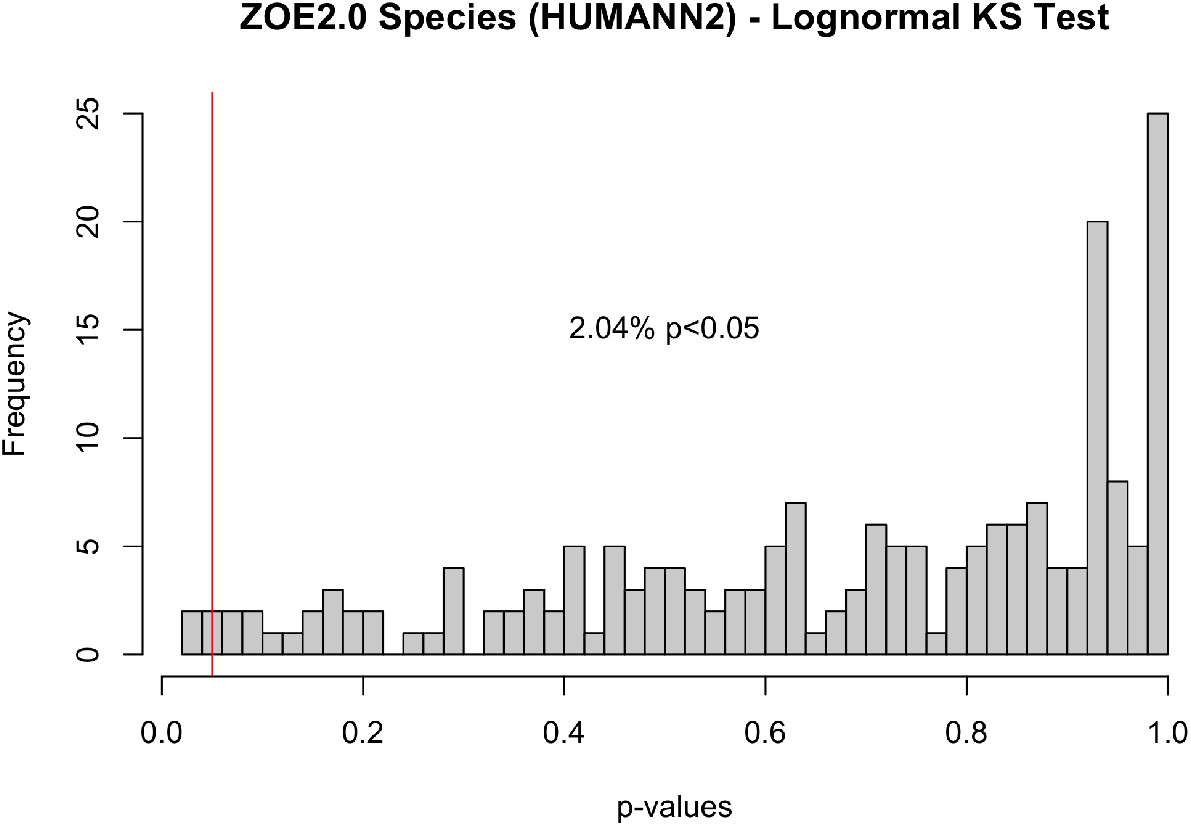
P-values obtained from lognormal (parameters from models fitted on nonzero counts for each metabolite and species) Kolmogorov-Smirnov test for ZOE 2.0 metabolites and Kraken2/Bracken microbiome species.

**Figure A3.**
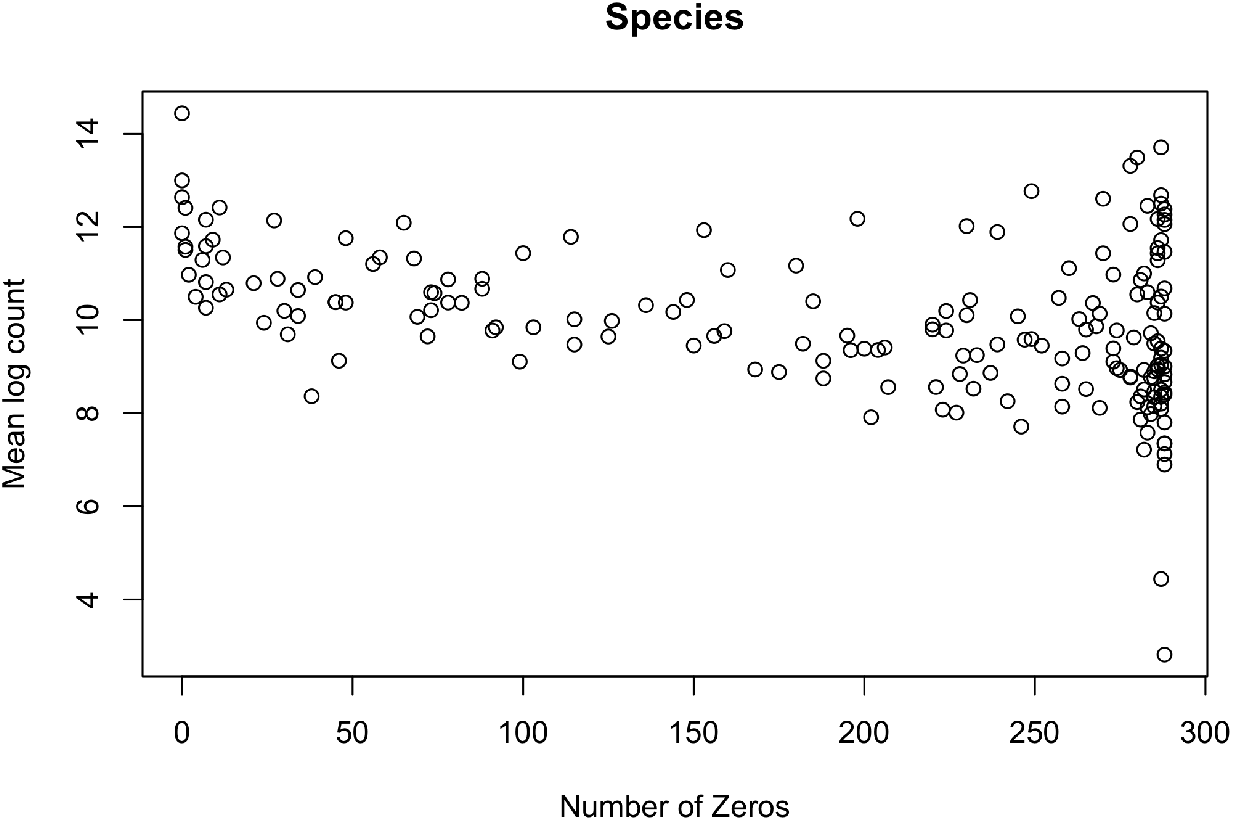
Species-wise (HUMAnN 2.0) numbers of zeros plotted against mean log nonzero counts.

**Figure A4.**
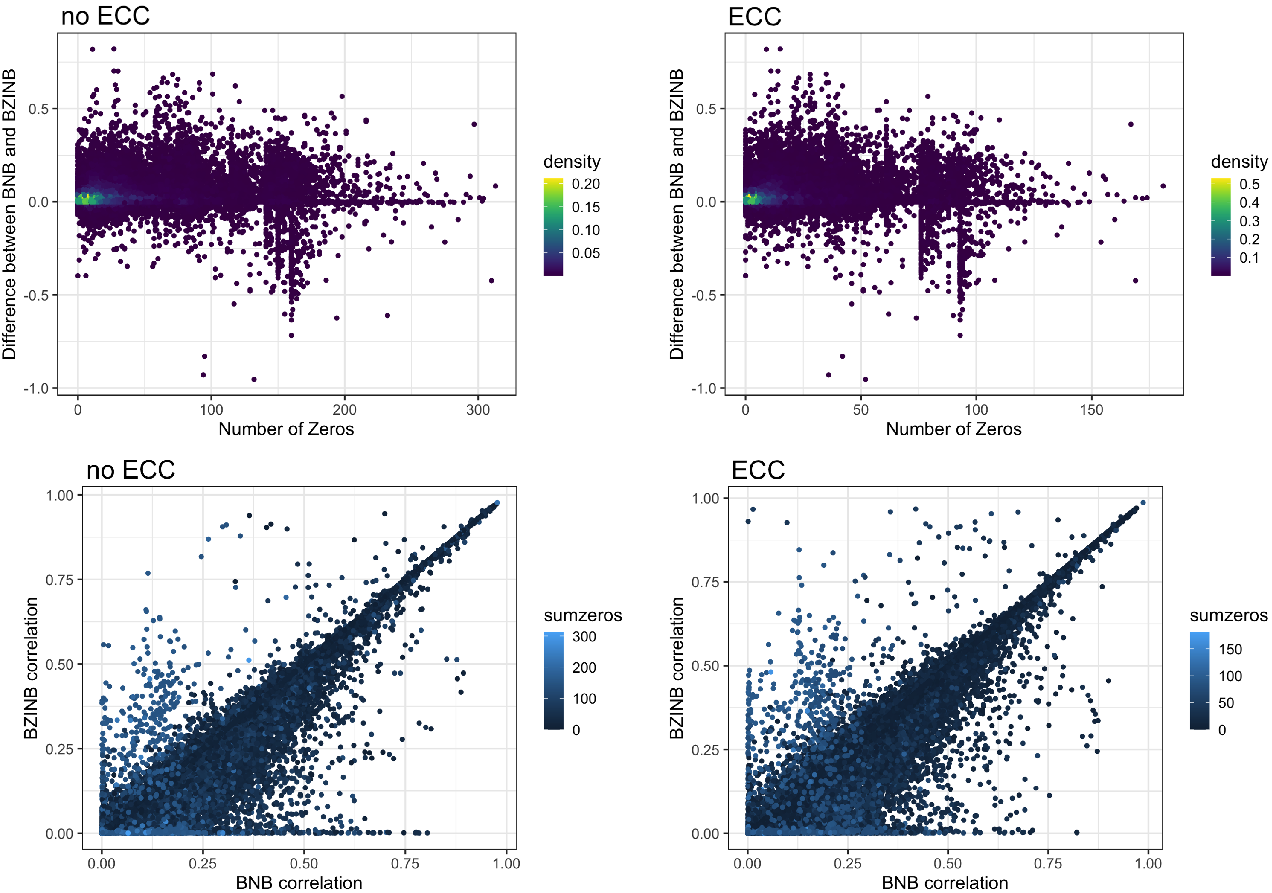
Comparison of BNB and BZINB correlations between all pairs of microbial species in ZOE2.0 with respect to the total number of zeros in each pair.

**Figure A5.**
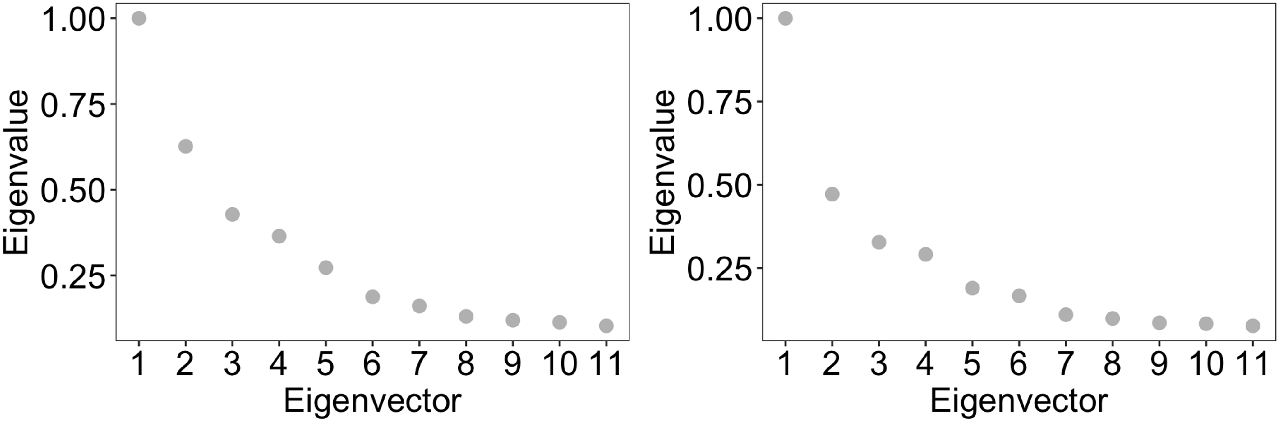
Eigenvalues of the Laplacian graph based on each affinity matrix for health and disease (ECC) groups in ZOE 2.0 Kraken2/Bracken microbiome data, which is used to determine an appropriate number of clusters.

**Figure A6:**
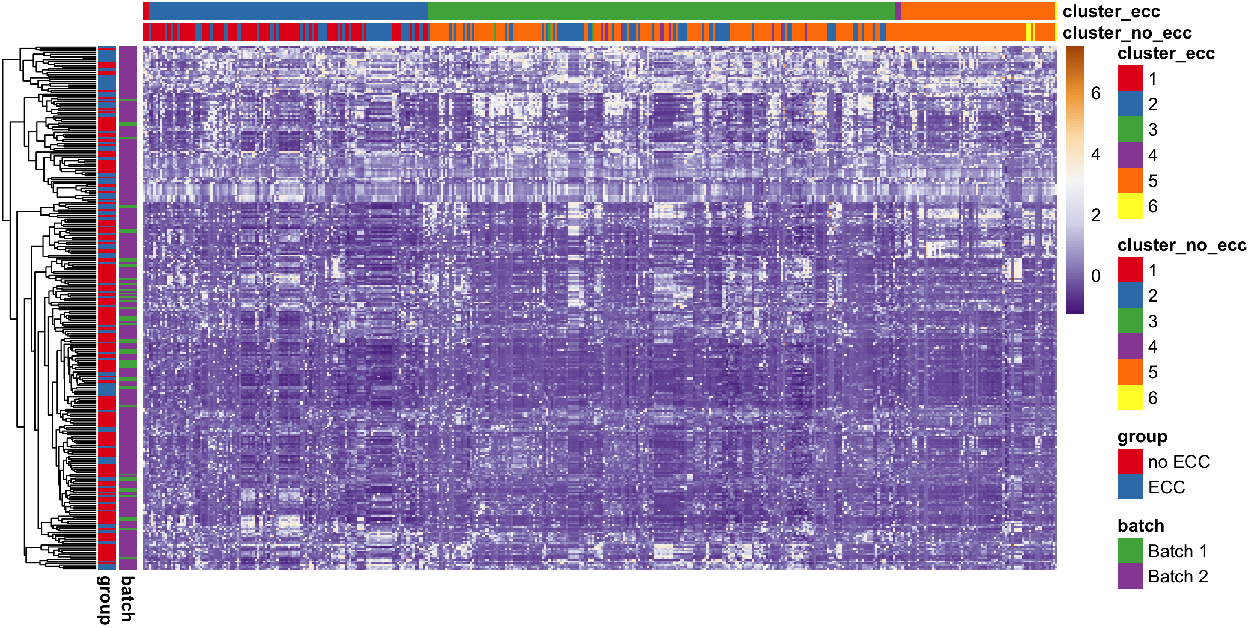
Heatmap of species abundance illustrating species modules identified by spectral clustering (shown as the two top bars). Columns represent individual species and are ordered by the clusters predicted from the affinity matrix based on the BZINB correlations between species in the diseased (ECC) group. Columns are annotated to show and compare the predicted clusters between health (no ECC) and disease (ECC) groups. Each row represents a participant (n=289). The rows are ordered based on hierarchical clustering and are annotated to illustrate health and disease groups and the sequencing batch. Standardized counts were used.

**Figure A7.**
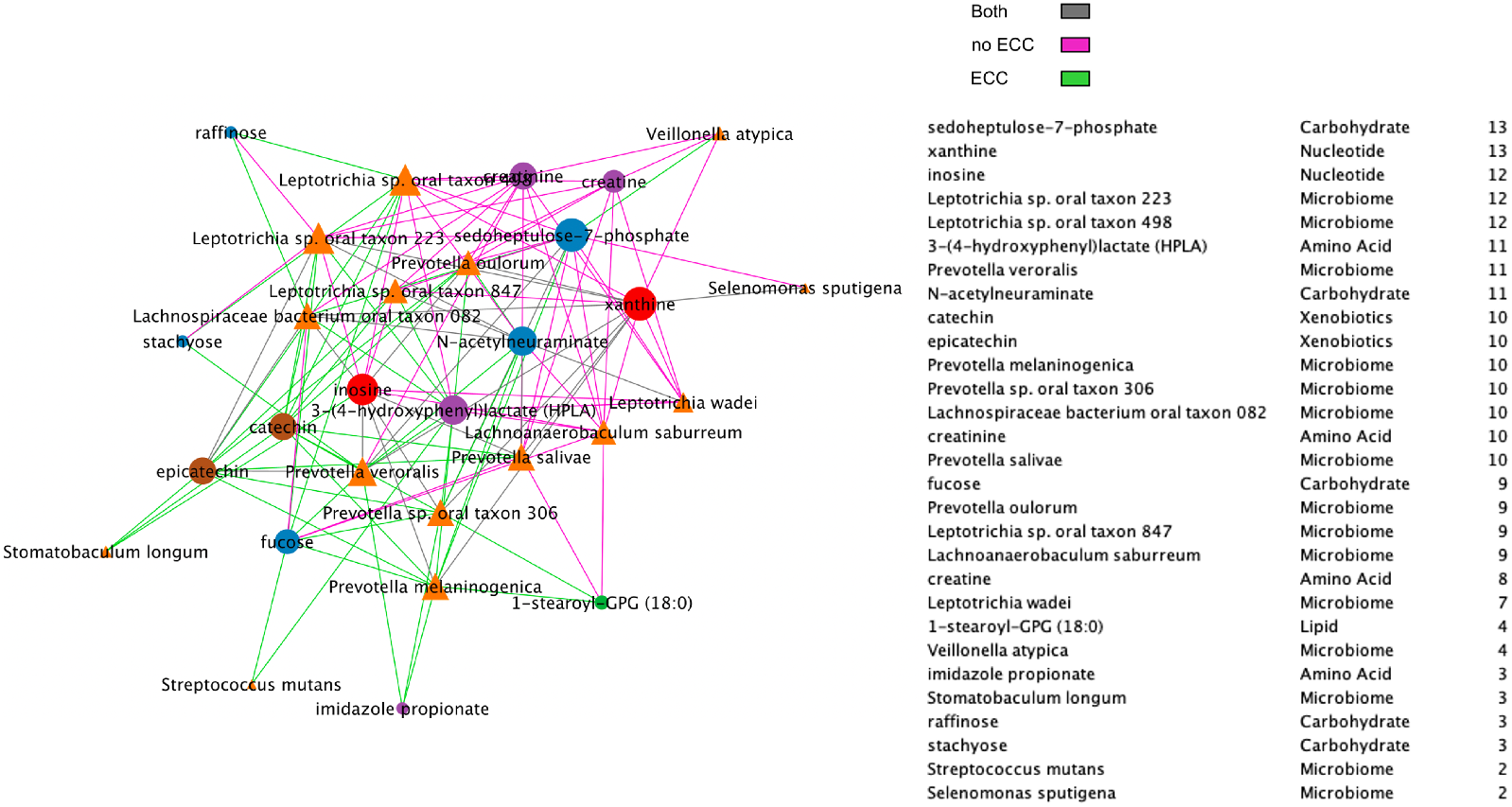
Spearman microbiome-metabolome correlation network including a node degree table. The strongest 30% absolute correlations are illustrated. Line colors represent correlations’ strength in health, disease (ECC), or both.

## References

1. Bauer, M. A.; Kainz, K.; Carmona-Gutierrez, D.; Madeo, F. Microbial Wars: Competition in Ecological Niches and within the Microbiome. Microbial Cell 2018, 5, 215–219, DOI:10.15698/mic2018.05.628.

2. Tong, H.; Chen, W.; Merritt, J.; Qi, F.; Shi, W.; Dong, X. Streptococcus Oligofermentans Inhibits Streptococcus Mutans through Conversion of Lactic Acid into Inhibitory H2O2: A Possible Counteroffensive Strategy for Interspecies Competition. Molecular Microbiology 2007, 63, doi:10.1111/j.1365-2958.2006.05546.x.

3. Nyvad, B.; Crielaard, W.; Mira, A.; Takahashi, N.; Beighton, D. Dental Caries from a Molecular Microbiological Perspective. Caries Research 2012, 47, 89–102, DOI:10.1159/000345367.

4. Mira, A.; Simon-Soro, A.; Curtis, M. A. Role of Microbial Communities in the Pathogenesis of Periodontal Diseases and Caries. Journal of Clinical Periodontology 2017, 44, S23–S38, DOI:10.1111/jcpe.12671.

5. Cho, H.; Ren, Z.; Divaris, K.; Roach, J.; Lin, B.; Lin, C.; Azcarate-Peril, A.; Simancas-Pallares, M.; Shrestha, P.; Orlenko, A.; Ginnis, J.; North, K.; Zandona, A. F.; Ribeiro, A.; Wu, D.; Koo, H. Pathobiont-Mediated Spatial Structuring Enhances Biofilm Virulence in Childhood Oral Disease. bioRxiv 2022, DOI:10.21203/rs.3.rs-1748651/v1.

6. Wu, N.; Yin, F.; Ou-Yang, L.; Zhu, Z.; Xie, W. Joint Learning of Multiple Gene Networks from Single-Cell Gene Expression Data. Computational and Structural Biotechnology Journal 2020, 18, 2583–2595, DOI:10.1016/j.csbj.2020.09.004.

7. Zhang, Z.; Zhang, X. Inference of High-Resolution Trajectories in Single-Cell RNA-Seq Data by Using RNA Velocity. Cell Reports Methods 2021, 1, 100095, DOI:10.1016/j.crmeth.2021.100095.

8. Gan, Y.; Liang, S.; Wei, Q.; Zou, G. Identification of Differential Gene Groups From Single-Cell Transcriptomes Using Network Entropy. Frontiers in Cell and Developmental Biology 2020, 8, doi:10.3389/fcell.2020.588041.

9. Ray, S.; Lall, S.; Bandyopadhyay, S. CODC: A Copula-Based Model to Identify Differential Coexpression. npj Systems Biology and Applications 2020, 6, doi:10.1038/s41540-020-0137-9.

10. Langfelder, P.; Horvath, S. WGCNA: An R Package for Weighted Correlation Network Analysis. BMC Bioinformatics 2008, 9, doi:10.1186/1471-2105-9-559.

11. Franzosa, E. A.; McIver, L. J.; Rahnavard, G.; Thompson, L. R.; Schirmer, M.; Weingart, G.; Lipson, K. S.; Knight, R.; Caporaso, J. G.; Segata, N.; Huttenhower, C. Species-Level Functional Profiling of Metagenomes and Metatranscriptomes. Nature Methods 2018, 15, 962–968, DOI:10.1038/s41592-018-0176-y.

12. Franzosa, E. A.; Sirota-Madi, A.; Avila-Pacheco, J.; Fornelos, N.; Haiser, H. J.; Reinker, S.; Vatanen, T.; Hall, A. B.; Mallick, H.; McIver, L. J.; Sauk, J. S.; Wilson, R. G.; Stevens, B. W.; Scott, J. M.; Pierce, K.; Deik, A. A.; Bullock, K.; Imhann, F.; Porter, J. A.; Zhernakova, A.; Fu, J.; Weersma, R. K.; Wijmenga, C.; Clish, C. B.; Vlamakis, H.; Huttenhower, C.; Xavier, R. J. Gut Microbiome Structure and Metabolic Activity in Inflammatory Bowel Disease. Nature Microbiology 2018, 4, 293–305, DOI:10.1038/s41564-018-0306-4.

13. Van Buren, E.; Hu, M.; Weng, C.; Jin, F.; Li, Y.; Wu, D.; Li, Y. TWO-SIGMA: A Novel Two-component Single Cell Model-based Association Method for Single-cell RNA-seq Data. Genetic Epidemiology 2020, 45, 142–153, DOI:10.1002/gepi.22361.

14. Cho, H.; Liu, C.; Preisser, J. S.; Wu, D. A bivariate zero-inflated negative binomial model and its applications to biomedical settings. bioRxiv 2020, DOI:10.1101/2020.03.06.977728.

15. Qiu, P. Embracing the Dropouts in Single-Cell RNA-Seq Analysis. Nature Communications, 2020, 11, doi:10.1038/s41467-020-14976-9.

16. Shi, J.; Malik, J. Normalized Cuts and Image Segmentation. IEEE Transactions on Pattern Analysis and Machine Intelligence 2000, 22, 888–905, DOI:10.1109/34.868688.

17. Wood, D. E.; Lu, J.; Langmead, B. Improved Metagenomic Analysis with Kraken 2. Genome Biology 2019, 20, doi:10.1186/s13059-019-1891-0.

18. Lu, J.; Breitwieser, F. P.; Thielen, P.; Salzberg, S. L. Bracken: Estimating Species Abundance in Metagenomics Data. PeerJ Computer Science 2017, 3, e104, DOI:10.7717/peerj-cs.104.

19. Dewhirst, F. E.; Chen, T.; Izard, J.; Paster, B. J.; Tanner, A. C. R.; Yu, W.-H.; Lakshmanan, A.; Wade, W. G. The Human Oral Microbiome. Journal of Bacteriology 2010, 192, 5002–5017, DOI:10.1128/jb.00542-10.

20. Cho, H.; Qu, Y.; Liu, C.; Tang, B.; Lyu, R.; Lin, B. M.; Roach, J.; Azcarate-Peril, M. A.; de Aguiar Ribeiro, A.; Love, M. I.; Divaris, K.; Wu, D. Comprehensive Evaluation of Methods for Differential Expression Analysis of Metatranscriptomics Data. bioRxiv 2021, DOI:10.1101/2021.07.14.452374.

21. Evans, A. M.; DeHaven, C. D.; Barrett, T.; Mitchell, M.; Milgram, E. Integrated, Nontargeted Ultrahigh Performance Liquid Chromatography/Electrospray Ionization Tandem Mass Spectrometry Platform for the Identification and Relative Quantification of the Small-Molecule Complement of Biological Systems. Analytical Chemistry 2009, 81, 6656–6667, DOI:10.1021/ac901536h.

22. Evans, A. M.; Bridgewater, B. R.; Liu, Q.; Mitchell, M. W.; Robinson, R. J.; Dai, H.; Stewart, S. J.; DeHaven, C. D.; Miller, L. A. D. High Resolution Mass Spectrometry Improves Data Quantity and Quality as Compared to Unit Mass Resolution Mass Spectrometry in High-Throughput Profiling Metabolomics. Journal of Postgenomics Drug Biomarker Development 2014, 4, doi:10.4172/2153-0769.1000132.

23. Divaris, K.; Slade, G. D.; Ferreira Zandona, A. G.; Preisser, J. S.; Ginnis, J.; Simancas-Pallares, M. A.; Agler, C. S.; Shrestha, P.; Karhade, D. S.; Ribeiro, A. de A.; Cho, H.; Gu, Y.; Meyer, B. D.; Joshi, A. R.; Azcarate-Peril, M. A.; Basta, P. V.; Wu, D.; North, K. E. Cohort Profile: ZOE 2.0—A Community-Based Genetic Epidemiologic Study of Early Childhood Oral Health. International Journal of Environmental Research and Public Health 2020, 17, 8056, DOI:10.3390/ijerph17218056.

24. Berahmand, K.; Nasiri, E.; Pir mohammadiani, R.; Li, Y. Spectral Clustering on Protein-Protein Interaction Networks via Constructing Affinity Matrix Using Attributed Graph Embedding. Computers in Biology and Medicine 2021, 138, 104933 DOI:10.1016/j.compbiomed.2021.104933.

25. Meilă, M.; Pentney, W. Clustering by weighted cuts in directed graphs. In Proceedings of the 2007 SIAM International Conference on Data Mining, Minneapolis, MN, USA, 26-28 April 2007.

26. John, C. R.; Watson, D.; Barnes, M. R.; Pitzalis, C.; Lewis, M. J. Spectrum: Fast Density-Aware Spectral Clustering for Single and Multi-Omic Data. Bioinformatics 2019, DOI:10.1093/bioinformatics/btz704.

27. Heimisdottir, L. H.; Lin, B. M.; Cho, H.; Orlenko, A.; Ribeiro, A. A.; Simon-Soro, A.; Roach, J.; Shungin, D.; Ginnis, J.; Simancas-Pallares, M. A.; Spangler, H. D.; Zandoná, A. G. F.; Wright, J. T.; Ramamoorthy, P.; Moore, J. H.; Koo, H.; Wu, D.; Divaris, K. Metabolomics Insights in Early Childhood Caries. Journal of Dental Research 2021, 100, 615–622, DOI:10.1177/0022034520982963.

28. Shannon, P.; Markiel, A.; Ozier, O.; Baliga, N. S.; Wang, J. T.; Ramage, D.; Amin, N.; Schwikowski, B.; Ideker, T. Cytoscape: A Software Environment for Integrated Models of Biomolecular Interaction Networks. Genome Research 2003, 13, 2498–2504, DOI:10.1101/gr.1239303.

29. Takahashi, N.; Washio, J.; Mayanagi, G. Metabolomic Approach to Oral Microbiota. Interface Oral Health Science 2011 2012, 334–340, DOI:10.1007/978-4-431-54070-0_98.

30. Takahashi, N. Microbial Ecosystem in the Oral Cavity: Metabolic Diversity in an Ecological Niche and Its Relationship with Oral Diseases. International Congress Series 2005, 1284, 103–112, DOI:10.1016/j.ics.2005.06.071.

31. Takahashi, N.; Washio, J.; Mayanagi, G. Metabolomic Approach to Oral Biofilm Characterization—A Future Direction of Biofilm Research. Journal of Oral Biosciences 2012, 54, 138–143, DOI:10.1016/j.job.2012.02.005.

32. Takahashi N. Oral Microbiome Metabolism: From “ Who Are They?” to “ What Are They Doing?”. Journal of Dental Research, 2015, 94, 1628–1637, DOI:10.1177/0022034515606045.

33. Mashima, I.; Nakazawa, F. Interaction between Streptococcus Spp. and Veillonella Tobetsuensis in the Early Stages of Oral Biofilm Formation. Journal of Bacteriology 2015, 197, 2104–2111, DOI:10.1128/JB.02512-14.

34. Sola-Penna, M. Metabolic Regulation by Lactate. IUBMB Life 2008, 60, 605–608, DOI:10.1002/iub.97.

35. Larrabee, M. G. Lactate Metabolism and Its Effects on Glucose Metabolism in an Excised Neural Tissue. Journal of Neurochemistry 2002, 64, 1734–1741, DOI:10.1046/j.1471-4159.1995.64041734.x.

36. Wu, D.; Smyth, G. K. Camera: A Competitive Gene Set Test Accounting for Inter-Gene Correlation. Nucleic Acids Research 2012, 40, e133–e133, DOI:10.1093/nar/gks461.

37. Van Buren, E.; Hu, M.; Cheng, L.; Wrobel, J.; Wilhelmsen, K.; Su, L.; Li, Y.; Wu, D. TWO-SIGMA-G: A New Competitive Gene Set Testing Framework for scRNA-Seq Data Accounting for Inter-Gene and Cell–Cell Correlation. Briefings in Bioinformatics 2022, 23, doi:10.1093/bib/bbac084. Version January 30, 2023 submitted to Microorganisms https://www.mdpi.com/journal/microorganisms

